# Conserved bacterial genes for biosynthesis of the algal morphogen thallusin span land and sea

**DOI:** 10.64898/2026.05.21.724671

**Authors:** Nicolai Kallscheuer, Johann F. Ulrich, Myriel Staack, Yaming Liu, Mrinal K. Das, Melanie Westphahl, Martin Sperfeld, Hermann Holbl, Jonathan Hammer, Raimund Nagel, Severin Sasso, Shinichi Sunagawa, Julie A. Z. Zedler, Hans-Dieter Arndt, Christine Beemelmanns, Christian Jogler, Thomas Wichard

**Affiliations:** Department of Microbial Interactions, Institute for Microbiology, Friedrich Schiller University Jena, Jena, Germany; Institute for Inorganic and Analytical Chemistry, Friedrich Schiller University Jena, Jena, Germany; Department Antiinfectives from Microbiota, Helmholtz Institute for Pharmaceutical Research Saarland (HIPS), Saarbrücken, Germany; Institute for Organic Chemistry and Macromolecular Chemistry, Friedrich Schiller University Jena, Jena, Germany; Institute of Microbiology, ETH Zurich, Zurich, Switzerland; Institute of Biology, Plant Physiology, Leipzig University, Leipzig, Germany; Synthetic Biology of Photosynthetic Organisms, Matthias Schleiden Institute for Genetics, Bioinformatics and Molecular Botany, Friedrich Schiller University Jena, Jena, Germany; Saarland University, Saarbrücken, Germany; Cluster of Excellence Balance of the Microverse, Friedrich Schiller University Jena, Jena, Germany

**Keywords:** algal-bacterial interactions, marine bacteria, *Bacteroidota*, *Planctomycetota*, *Actinomycetota*, *Pseudomonadota*

## Abstract

Bacterial signals control the development of marine algae, yet the molecular basis of these cross-kingdom interactions remains largely unknown. Thallusin is the paradigmatic case: isolated in 2005, it induces rhizoid and cell wall formation in the green seaweed *Ulva* at picomolar concentrations, but its biosynthesis has remained elusive for two decades. Comparative genomics across five bacterial phyla identifies a conserved set of genes - the eustigmatophyte bacterial operon (*ebo*) - as determinants of thallusin biosynthesis. Isotope labeling, heterologous expression, and gene deletion in *Stieleria maiorica* show that the aromatic scaffold derives from a cyclitol precursor and L-aspartate, with subsequent prenylation and cyclization. Searching 124,295 prokaryotic genomes identifies producers in eleven bacterial lineages, including soil cyanobacteria, establishing thallusin as a widespread cross-kingdom signal reaching beyond the ocean.

## Introduction

Microbial metabolites shape the development, physiology, and ecology of multicellular eukaryotic hosts across the tree of life. Marine macroalgae host complex bacterial communities whose metabolites direct algal growth and morphogenesis. A striking case of such cross-kingdom signaling is thallusin, a potent bacterial morphogen governing development of the green macroalga *Ulva* (*1, 2*).

The meroterpenoid thallusin was isolated in 2005 from the marine bacterium *Cytophaga* sp. YM2-23 (*Bacteroidota*) (*1*). Low natural yields drove total synthesis of thallusin and analogs (*3–6*). This synthesis enabled structure-activity studies that defined the role of thallusin in bacterial control of *Ulva* morphogenesis. Axenic *Ulva* develops as undifferentiated, callus-like aggregates lacking rhizoids and proper cell walls (*7*). Co-culture with *Roseovarius* sp. and *Maribacter* sp., restores normal thallus development through bacterially produced growth- and morphogenesis-promoting factors. Within this community, *Roseovarius* sp. secretes an uncharacterized, cytokinin-like signal that drives cell division. *Maribacter-*derived thallusin acts as an auxin mimic, inducing rhizoid differentiation and organized cell wall formation (*7*) (Fig. S1, Table S1). Thallusin achieves these effects at picomolar concentrations (∼10⁻^11^ mol/L) in axenic gametes (*8*). Yet classical plant hormones - including indole-3-acetic acid, zeatin, and abscisic acid - cannot substitute (*9*), defining thallusin as a distinct marine morphogen. Thallusin also forms stable Fe(III) complexes, selectively imported by *Ulva* gametes (*6, 8*). This dual function - as morphogen and iron carrier - defines metabolite-mediated bacterial-algal symbiosis.

Until now, all reported thallusin producers belong to the phylum *Bacteroidota* (*8, 10*). Yet thallusin’s global ecological significance suggests biosynthesis across additional bacterial phyla in *Ulva*-associated communities (*11, 12*). Two decades after its structure was solved, the biosynthetic pathway remains unknown, despite the proposal of saccharoquinoline (from *Saccharomonospora* sp. CNQ-490) as precursor (*13*).

To address these gaps, we used selective cultivation to isolate slow growing, nutritionally specialized strains from underexplored *Ulva* microbiomes. Comparative genomics of these isolates and known producers and non-producers identify a conserved set of biosynthesis genes. Within this set, prenyltransferase-encoding genes - required for the formation of thallusin’s terpene moiety - guided chemical synthesis of pathway intermediates and isotopic precursor feeding performed in this study. We resolve thallusin’s biosynthetic origin, expanded producer diversity across multiple phyla, and show that phylum-specific gene organization enables reliable cultivation-independent identification of producers from metagenomes.

## Results and discussion

### Phylogenetically diverse bacteria produce the *Ulva* morphogen thallusin

Building on the hypothesis that saccharoquinoline serves as a biosynthetic precursor of thallusin (*13*), we investigated whether the actinomycetal strain *Saccharomonospora* sp. CNQ-490 - a previously reported saccharoquinoline producer - also produces thallusin. Applying our standardized analytical protocol, we detected thallusin in the spent cultivation medium of this strain, indicating that thallusin biosynthesis exceeds beyond *Bacteroidota* and may be widespread among phylogenetically diverse bacteria.

This discovery prompted a broader investigation of phylogenetically diverse *Ulva*-associated bacteria for thallusin production. For this, *Ulva* sp. specimens were collected from Heligoland Island (Germany, North Sea) and incubated using our previously described deep-cultivation procedure (*11, 12, 14*), yielding three novel isolates: *Rhodopirellula* sp. UH5 (*Planctomycetota*), *Maribacter* sp. UH1 and *Aurantibacter* sp. UH7 (both *Bacteroidota*). Thallusin production was assessed using two independent methods - LC-MS^2^ detection and an *Ulva*-based bioactivity assay - confirming thallusin production in strains UH1 and UH5, but not in strain UH7. The analysis was extended to 31 additional strains spanning multiple phyla, encompassing all previously confirmed *Bacteroidota* producers alongside planctomycetal isolates from a prior sampling expedition (*6, 10, 11, 15–18*) (Fig. 1A, Fig. S2, Table S2). Across all 35 strains examined, 27 were identified as thallusin producers and eight as non-producers, confirming thallusin biosynthesis across the environmentally relevant phyla *Bacteroidota*, *Actinomycetota, Pseudomonadota* and *Planctomycetota*.

**Fig. 1.**
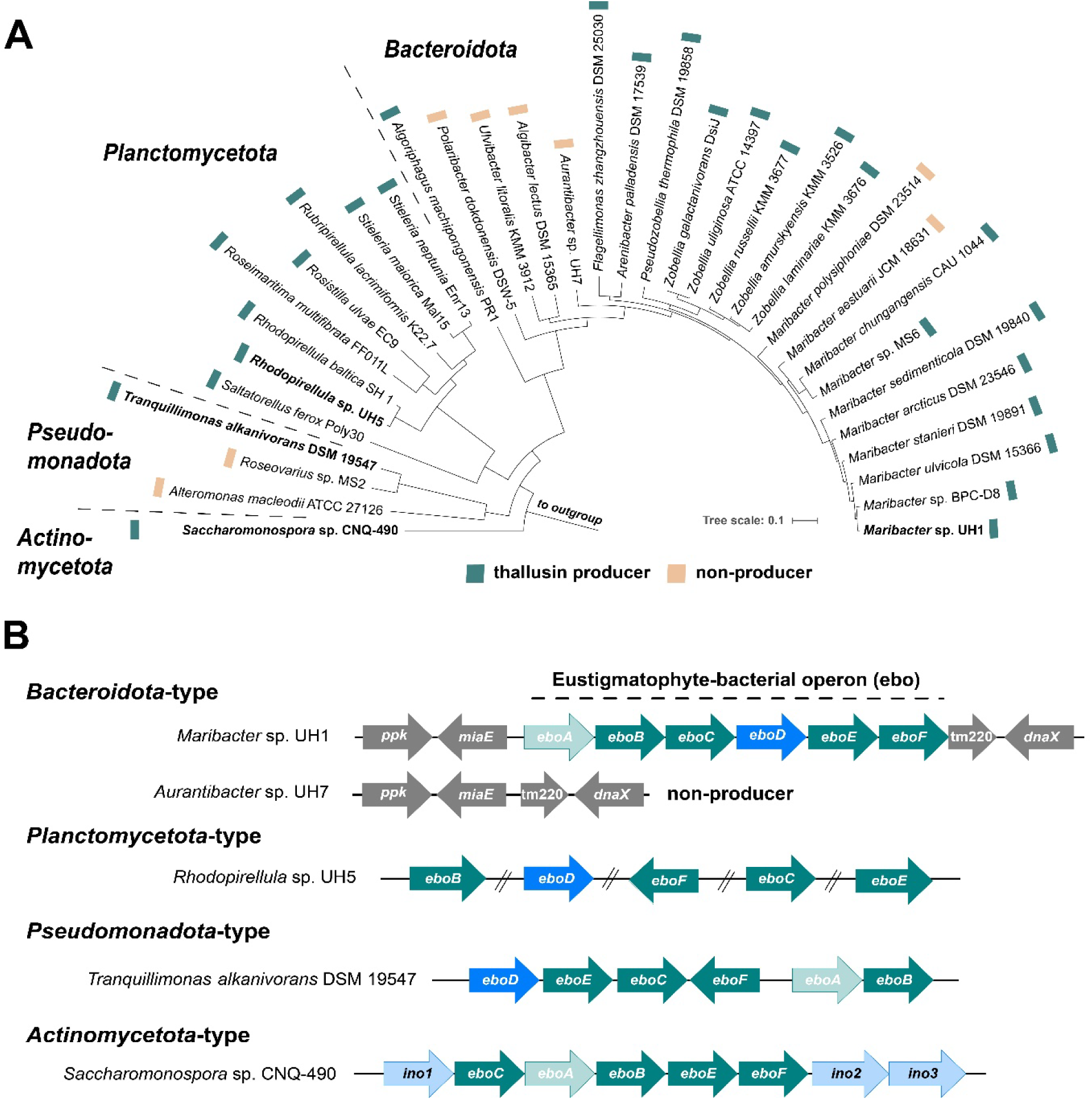
Comparison of experimentally confirmed thallusin producer and non-producer strains across four bacterial phyla. (**A**) Multi-locus sequence analysis-based maximum likelihood dendrogram visualizing the phylogenetic position of experimentally confirmed thallusin producers and non-producers. (**B**) Phylum-type organization of *ebo* genes involved in thallusin biosynthesis. A detailed comparison of clusters of all producers is provided in Fig. S5. NCBI RefSeq accession numbers of the analyzed genomes are provided in Table S2.

### Comparative genomics links thallusin production to the *ebo* genes

The occurrence of thallusin producers across phylogenetically diverse bacterial lineages provided an entry for identifying the underlying biosynthetic genes by applying comparative genomics. Candidate genes were evaluated using multiple approaches, including genome mining for biosynthetic gene clusters, re-annotation based on precomputed orthologous groups, and pangenome reconstruction with subtraction of non-producer genes from the minimal producer gene set. The most conclusive results emerged from the pangenome analysis of nine *Maribacter* strains - eight producers and one non-producer (Fig. S3). Of approximately 9,200 gene clusters in the pangenome, 91 were conserved exclusively across all producers (Table S3). Within this set, a search for prenyltransferase-encoding genes - candidates for terpenoid moiety formation characteristic for thallusin - identified a single gene encoding the UbiA-like prenyltransferase EboC (*19*). Inspection of its genomic context revealed that *eboC* is part of the six-gene operon *eboABCDEF* (*20*). The *ebo* (eustigmatophyte-bacterial operon) genes were named after eustigmatophyte algae, the proposed origin of the operon, which symbiotic bacteria may have acquired via horizontal gene transfer (*21*). The encoded enzymes have been proposed previously to produce an unknown prenylated cyclitol derivative (*21*), and are currently annotated as uncharacterized protein (EboA), TatD-like hydrolase (EboB), UbiA-like prenyltransferase (EboC), 3-dehydroquinate synthase-like sugar phosphate cyclase (EboD), xylose isomerase-like protein (EboE) and alkaline phosphatase family protein (EboE). BLAST analyses confirmed the presence of the *ebo* genes in all confirmed thallusin producer genomes across all phyla examined. They were consistently absent from all non-producers.

Despite the clear genotype-phenotype correlation, the link to thallusin biosynthesis was unexpected, as the *ebo* genes had previously been associated with precursor transport during biosynthesis of the sunscreen pigment scytonemin in the cyanobacterium *Nostoc punctiforme* (*22*). However, since the Ebo proteins are not directly involved in scytonemin biosynthesis, those observations may result from indirect effects (*22*). Given the ambiguous functional assignment in *N. punctiforme*, three cyanobacteria strains - two harboring and one lacking the *ebo* genes, including *N. punctiforme* itself - were tested for thallusin production. This was of particular interest because thallusin has thus far been reported to be produced exclusively by marine bacteria, whereas *N. punctiforme* is primarily terrestrial. Only the two *ebo* gene-harboring cyanobacterial strains produce thallusin, further reinforcing the functional link between the *ebo* genes and thallusin biosynthesis (Fig. S4).

Synteny analyses of the genes in confirmed thallusin producers revealed phylum-specific organizational patterns. Thallusin-producing *Bacteroidota* harbor the complete six-gene operon *eboABCDEF*, whereas the *Actinomycetota*- and *Pseudomonadota*-type arrangements differ in gene order (Fig. 1B). The *Actinomycetota*-type organization additionally lacks *eboD*. Instead, the operon is flanked on both sides by genes encoding enzymes putatively involved in inositol biosynthesis and conversion (*ino1*, *ino2* and *ino3*) (Fig. S5). In *Planctomycetota*, individual *ebo* genes occur without apparent clustering, distributed throughout the genomes, and *eboA* is absent (Fig. 1B). This scattered arrangement is consistent with the comparatively low frequency of operon structures in planctomycetes relative to other bacterial phyla (*23*).

To assess the global distribution and gene cluster architecture of *ebo* clusters, 124,295 prokaryotic, species-level representative genomes were screened for the presence and genomic co-localization of the six *ebo* and three *ino* genes. This identified 3,403 clusters in 3,304 genomes from eleven bacterial and two archaeal phyla (Fig. S6). The analysis confirmed the phylum-specific variations in gene organization, including the clustered versus isolated arrangement, and the absence of *eboD* or *eboA* in distinct phyla. In total, 4,405 genomes harbor the conserved core genes *eboB*, *eboC*, *eboE* and *eboF* that were frequently co-localized, supporting their function as a cohesive biosynthetic unit (Fig. S6).

### Stable isotope tracing identifies L-aspartate as a substrate for thallusin biosynthesis

Having established a founded hypothesis of thallusin biosynthesis, we next sought to identify the biosynthetic precursor metabolites. Retrobiosynthetic analysis of the meroterpenoid backbone of thallusin leads to a heteroaromatic starter unit and a cyclized drimane-type terpenoid moiety. The positions of the carboxyl group and the heterocyclic nitrogen atom within the nicotinic acid moiety suggested an α-amino acid as the biosynthetic precursor. To test for this possibility, stable isotope (^13^C, ^15^N) tracing experiments were conducted using the model producer *Maribacter* sp. MS6 that was cultivated in the presence of isotopically labeled amino acids and related precursors.

Isotope labeling experiments revealed that L-aspartate is efficiently incorporated into thallusin, as evident by strong enrichment and clear mass shifts upon feeding with ¹⁵N- and ¹³C-labeled aspartate (Fig. 3, Fig. S7A). MS^2^ analysis of the diagnostic quinoline-associated fragment ion of thallusin confirmed enrichment of all labeled atoms from ^15^N,^13^C_4_-aspartate, establishing L-aspartate as an entry metabolite for thallusin biosynthesis (Fig. 3C).

In contrast, ^13^C₁₁^15^N₂-tryptophan, ^13^C-erythrose, or ^15^N-anthranilic acid showed only minimal or no isotope incorporation (Fig. S7B), indicating that these building blocks are not directly involved in thallusin biosynthesis. Thus, thallusin biosynthesis does not proceed via the canonical shikimate/tryptophan pathway, despite structural similarity of its quinoline moiety to typical aromatic metabolites. However, feeding with ^15^N,^13^C_5_-glutamate resulted in partial labeling of both saccharoquinoline and thallusin (Fig. S7C), suggesting indirect incorporation via central metabolic interconversions. Among all metabolites tested, only L-aspartate yielded strong and reproducible isotope incorporation, indicating direct and complete incorporation into thallusin.

### EboC catalyses the farnesylation of hydroxylated quinolines

To validate EboC as a prenyltransferase, synthetic codon-optimized genes encoding EboC from *Maribacter* sp. MS6 (*Bacteroidota*), *Stieleria neptunia* Enr13 (*Planctomycetota*) and *Saccharomonospora* sp. CNQ-490 (*Actinomycetota*) were heterologously expressed in *Escherichia coli*. Membrane fractions containing EboC were subjected to enzymatic assays with farnesyl pyrophosphate (FPP) and aromatic substrates. All three proteins accepted hydroxylated quinoline derivatives as substrates, with FPP serving as the prenyl donor (Fig. S8). Among the tested quinaldic acid substrates, 6,7,8-trihydroxyquinaldic acid (6,7,8-HQA) ethyl ester yielded the highest conversion to the prenylated product for all three enzymes as assessed by the ion signal intensity of the farnesylated derivative.

### Deletion of individual *ebo* core genes abolishes thallusin biosynthesis

To determine whether each *ebo* core gene (*eboB, C*, *E* and *F*) is individually required for thallusin biosynthesis, targeted single-gene deletion mutants were constructed. A strain in which the *ebo* genes are scattered throughout the genome was selected to avoid potential polar effects on gene expression arising from deletions within an operon. *Stieleria maiorica* Mal15, a genetically tractable planctomycetal thallusin producer, was used for this purpose. UHPLC-ESI-HRMS analysis confirmed that all four deletion mutants (Δ*eboB, C*, *E* and *F*) completely lost the ability to synthesize thallusin, consistent with results from the *Ulva* morphogenesis bioassays (Fig. 2). The wild-type strain restored the complete algal morphotype in the presence of *Roseovarius* sp. MS2, characterized by a well-organized thallus and distinct rhizoid formation (type IV, Table S1). In contrast, all deletion mutants induced only the incomplete morphotype characteristic of non-producers, yielding *Ulva* phenotypes with surface protrusions, absence of rhizoids and defined cell walls (type II, Table S1) and thus reduced growth (Fig. S9). Collectively, these results demonstrate that each *ebo* core gene is essential for thallusin biosynthesis and the associated morphogenetic activity.

**Fig. 2.**
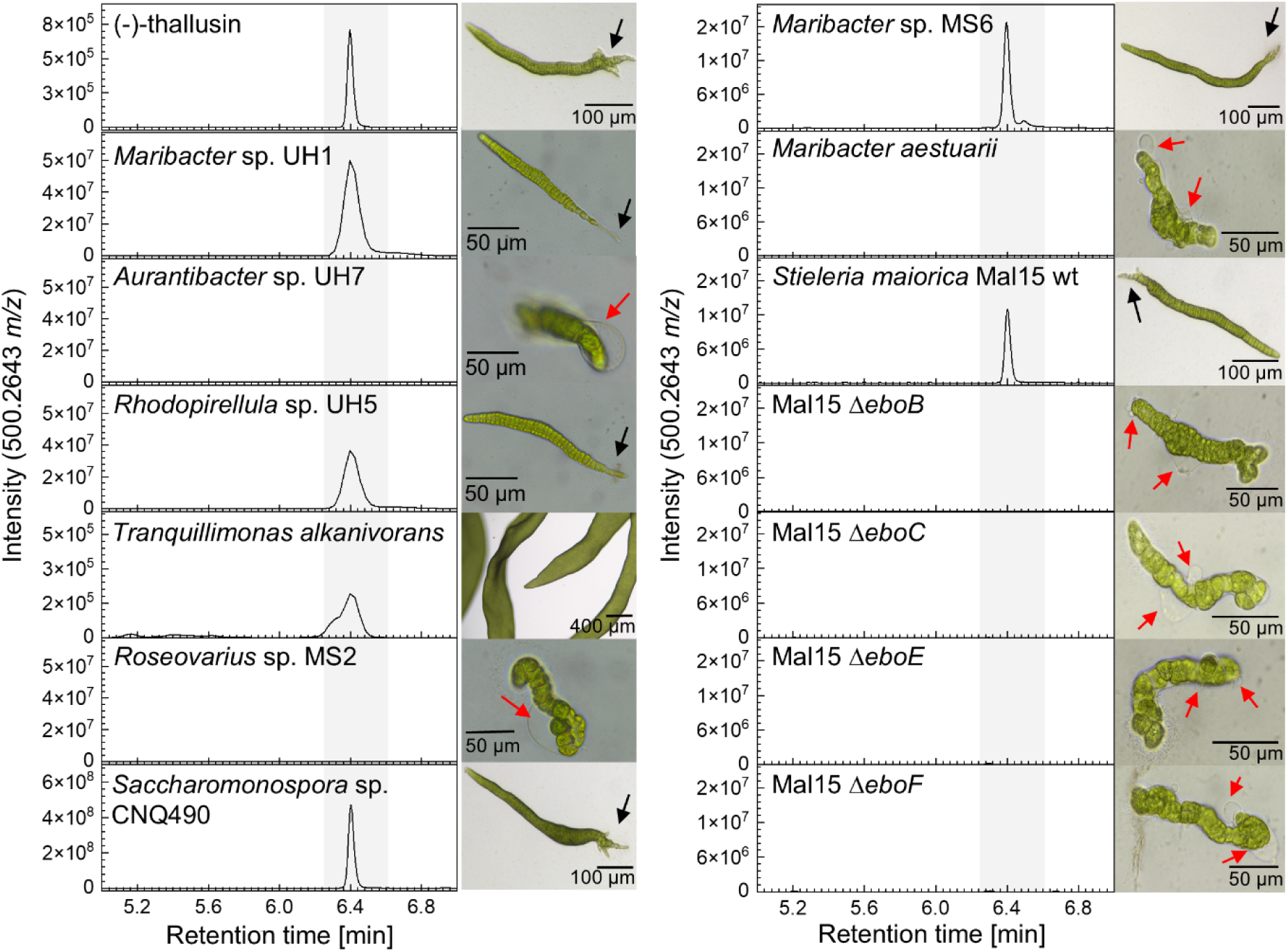
Thallusin production across bacterial phyla and functional validation of the *ebo* genes. Extracted ion chromatograms (EIC; *m/z* 500.2643) of derivatized thallusin in bacterial cultures. Thallusin-producing strains (*Maribacter* sp. UH1, *Rhodopirellula* sp. UH5, *Maribacter* sp. MS6, *Saccharomonospora* sp. CNQ490) show a characteristic peak at ∼6.4 min, whereas non-producers (*Aurantibacter* sp. UH7, *Maribacter aestuarii*, *Roseovarius* sp. MS2) lack the signal. Corresponding *Ulva* morphogenesis bioassays were performed in the presence of *Roseovarius* sp. MS2 and the potential thallusin producers. Thallusin-producing strains restore normal *Ulva* morphology with elongated thalli and rhizoids (black arrows), while non-producers induce aberrant morphotypes with cell aggregates and protrusions (red arrows) in the presence of *Roseovarius* sp. MS2. Deletion of *ebo* genes in *Stieleria maiorica* Mal15 abolishes thallusin production and results in thallusin-deficient phenotypes, whereas the wild type restores full morphogenesis. Scale bars, 50-400 µm.

**Fig. 3.**
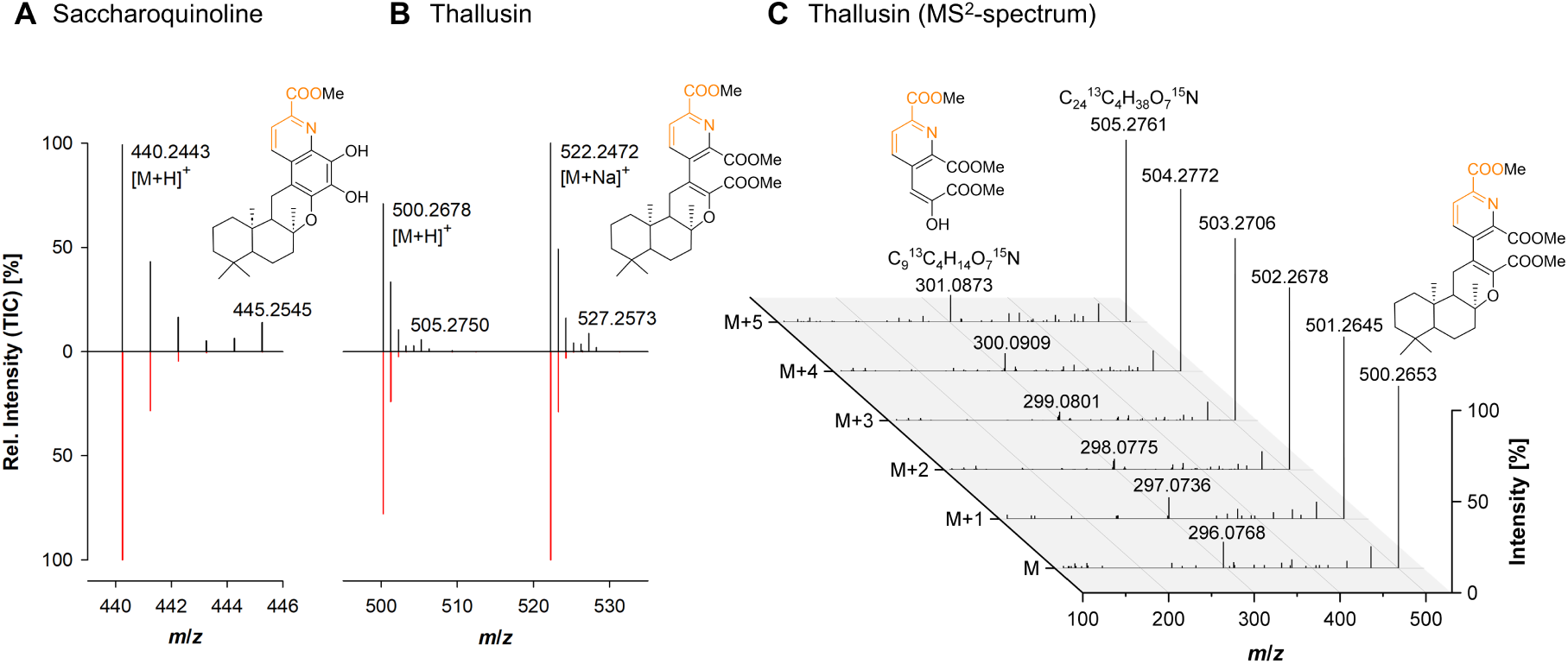
Isotope tracing reveals incorporation of aspartate into thallusin. Mass spectra of monomethylated saccharoquinoline (**A**) and fully methylated thallusin (**B**). Molecular structures are shown in their neutral form [M]. Feeding with ¹⁵N,¹³C_4_-aspartate leads to incorporation into both the monomethylated derivative of saccharoquinoline (*m/z* 440.2443) and the fully methylated thallusin (*m/z* 500.2678 [M+H]⁺, 522.2472 [M+Na]⁺), demonstrating that aspartate contributes directly to thallusin biosynthesis. The isotopic signature of the mass spectra corresponds to the unlabeled compound (red) and to the isotopically labeled compound derived from feeding experiments (black) (**C**) MS² spectra of this derivatized and labeled thallusin reveal a stepwise isotopologue pattern (M to M+5). The diagnostic quinoline-derived fragment (*m/z* ∼296-301) exhibits corresponding mass shifts, demonstrating incorporation of all four ¹³C atoms and one ¹⁵N atom from aspartate into the non-terpenoid quinoline core of thallusin (orange).

### Biosynthetic considerations for thallusin production

Our analyses establish a shared biosynthetic relationship between saccharoquinoline and thallusin and identify the *ebo* genes as the conserved genetic basis underlying this pathway across phylogenetically diverse bacteria (Fig. 4). The consistent co-occurrence of thallusin production with *ebo* core genes across all experimentally validated producer strains, together with the detection of saccharoquinoline, establishes the genetic and biochemical basis of thallusin biosynthesis. While the organization of the *ebo* loci varies substantially between bacterial phyla, a conserved set of core biosynthetic genes is retained in all producers, indicating a shared metabolic logic despite lineage-specific adaptations. Besides *eboA*, which could not be identified in most *Planctomycetota* genomes (*20*), and *eboD*, which is absent in *Actinomycetota*, the remaining core genes are consistently conserved across all validated producers, supporting their direct involvement in thallusin formation.

**Fig. 4.**
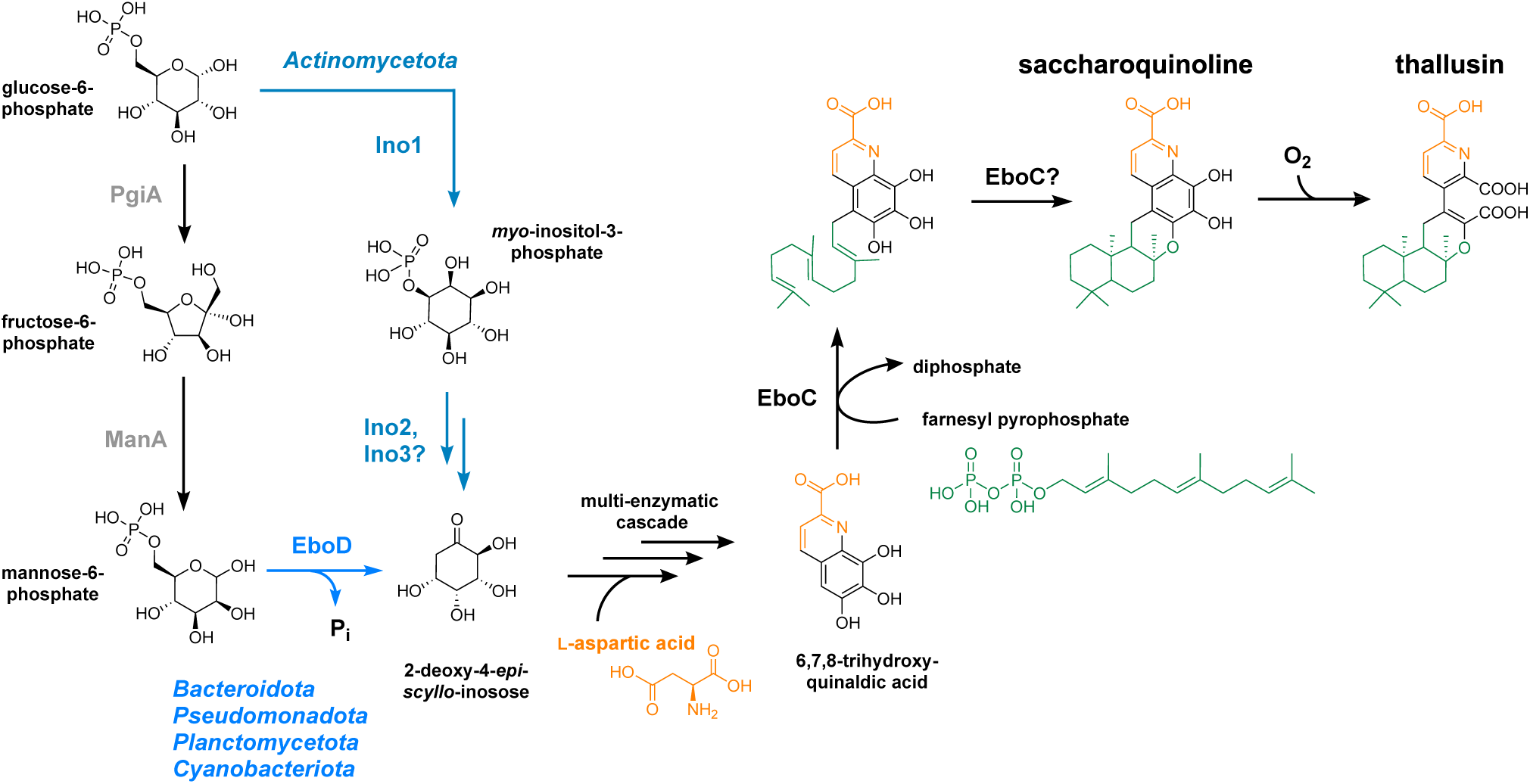
Postulated thallusin biosynthetic route. The biosynthesis of thallusin starts from 2-deoxy-4-*epi*-*scyllo*-inosose that can be produced by two different pathways starting from glucose-6-phosphate, either via cyclization of mannose-6-P catalyzed by EboD (in all strains that harbor *eboD*) or via a bypass route probably with inositol-3-P and two unknown compounds as pathway intermediates (in *Actinomycetota*).

Our comparative genomic and isotope-labeling analyses now link thallusin biosynthesis to a cyclitol as precursor. EboD was recently characterized as a member of a novel class of sugar phosphate cyclases converting mannose-6-phosphate into 2-deoxy-4-*epi*-*scyllo*-inosose (*24*). The enzyme is encoded in thallusin-producing members of the *Bacteroidota*, *Planctomycetota*, *Pseudomonadota*, and *Cyanobacteriota* (Fig. S6) and may catalyze the initial and committed step towards thallusin biosynthesis. While in *Actinomycetota eboD* homologs were not detectable, the co-occurrence with genes of inositol metabolism, including inositol-3-phosphate synthase (Ino1) and two putative isomerases/dehydratases (Ino2 and Ino3), is suggesting an alternative route from glucose-6-phosphate instead of mannose-6-phosphate via inositol-derived intermediates that could converge on the same precursor structure (Fig. 4). Together with the experimentally verified incorporation of aspartate into the quinoline moiety, these findings establish the first biosynthetic model for the formation of the non-terpenoid core of thallusin and enable the mechanistic investigation of this long-elusive pathway.

The UbiA-like prenyltransferase EboC catalyzed the farnesylation of 6,7,8-HQA ethyl ester in homologs derived from phylogenetically distinct thallusin producers, establishing prenylation of hydroxylated quinoline intermediates as a conserved step in thallusin biosynthesis (*19*). These findings experimentally connect the aromatic precursor pathway to assembly of the terpenoid moiety and provide biochemical evidence for the proposed pathway intermediate. The drimane-type sesquiterpene residue of thallusin is structurally analogous to that found in the meroterpenoid BE-40644 from *Actinoplanes* sp. A40644 (*25*). Notably, the BE-40644 biosynthetic gene cluster lacks a dedicated terpene cyclase yet remains functional upon heterologous expression (*26*), suggesting that cyclization can occur in the absence of a dedicated cyclase. Together with recent evidence implicating UbiA-like prenyltransferases in terpene cyclization reactions (*27*), our results raise the possibility that EboC catalyzes both prenylation and subsequent cyclization during thallusin assembly, thereby identifying EboC as a central catalyst of late-stage thallusin biosynthesis.

One of the remaining questions in thallusin biosynthesis is the biochemical basis of the oxidative rearrangement converting saccharoquinoline into thallusin. This transformation is structurally reminiscent of the dioxygen-dependent intradiol ring-cleavage of hydroxybenzoates, such as catechol and protocatechuic acid, that initiates aerobic aromatic compound degradation via the β-ketoadipate pathway (*28*). However, neither the Ebo proteins nor comparative genome mining identified conserved Fe³⁺-dependent intradiol dioxygenases or other obvious ring-cleavage enzymes associated with the pathway. The identification of saccharoquinoline and thallusin production within the same biosynthetic context now enables targeted investigation of these late-stage conversion reactions. Given the metal-chelating properties of thallusin, oxidative processes involving iron-assisted rearrangements or other non-canonical oxidative chemistry may contribute to the formation of the characteristic ring-opened scaffold.

### Mapping thallusin biosynthetic potential across the prokaryotic tree of life

Until now, thallusin biosynthesis had been associated exclusively with marine members of the phylum *Bacteroidota*. Here, the identification of thallusin production in *Actinomycetota*, *Pseudomonadota*, *Planctomycetota*, and *Cyanobacteriota*, together with the biosynthetic link between saccharoquinoline and thallusin, substantially expands the phylogenetic and metabolic framework of thallusin biosynthesis. Extending this analysis to 124,295 representative prokaryotic genomes revealed the conserved *ebo* core genes in at least six additional bacterial phyla, including *Spirochaetota*, *Deinococcota*, *Verrucomicrobiota, Acidobacteriota, Myxococcota, and Desulfobacterota* (Fig. 5), demonstrating that the genetic potential for thallusin biosynthesis is far more widespread than previously recognized.

**Fig. 5.**
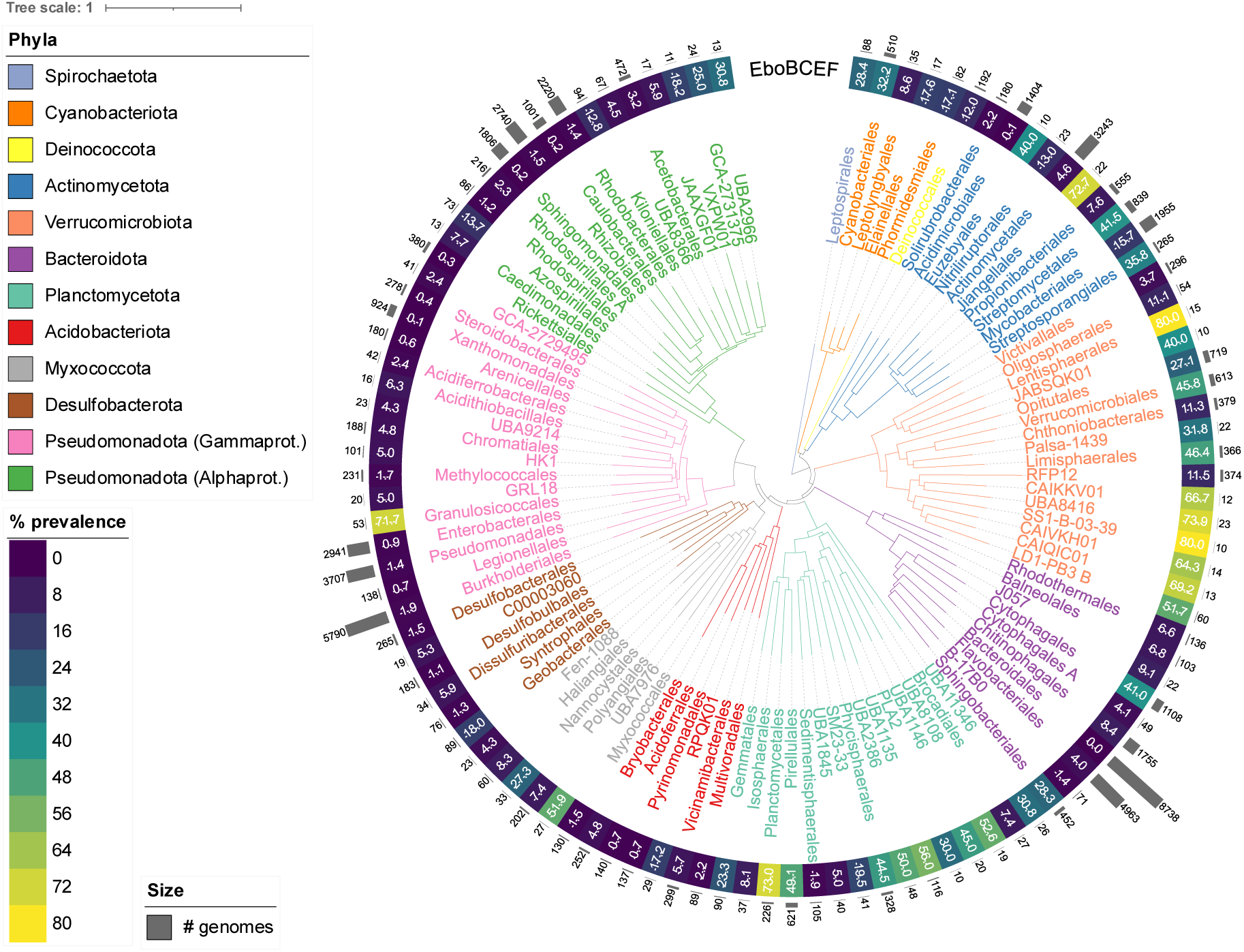
Mapping the prevalence of thallusin biosynthetic genes across the bacterial tree of life. The tree depicts the combined prevalence of the identified core genes *eboB*, *eboC*, *eboE* and *eboF* genes (indicated by the heatmap ring) overlayed onto the GTDB bacterial taxonomic tree, collapsed at the order level. The tree was filtered to include only the 11 bacterial phyla with robust thallusin biosynthesis potential, excluding orders represented by fewer than ten genomes (indicated by outer bar-plot ring), and retaining only orders in which at least one genome contains all four core genes.

The distribution of the *ebo* core genes was uneven across bacterial diversity. While the gene set occurred across multiple orders within *Actinomycetota*, *Verrucomicrobiota*, and *Planctomycetota*, it was restricted to specialized lineages in other phyla, such as the order *Granulosicoccales* within the *Pseudomonadota*. Members of *Granulosicoccales*, including the genus *Granulosicoccus*, are recurrent constituents of macroalgal surface microbiomes, including those associated with *Ulva* (*29, 30*), similar to members of the *Verrucomicrobiota* (*11, 31*). This distribution suggests that these taxa may contribute to thallusin-mediated algal morphogenesis, although their precise ecological roles remain unresolved.

Notably, the occurrence of *ebo* genes in characteristic soil- and freshwater-associated taxa, including *Nostoc* spp., extends the potential ecological range of thallusin producers far beyond marine environments and suggests that thallusin-mediated signaling may operate across diverse terrestrial and aquatic ecosystems.

## Conclusion

Identification of the *ebo* genes as the biosynthetic determinants of thallusin resolves a two-decade-old question in bacterial–algal chemical communication and enables cultivation-independent identification of thallusin-producing microorganisms and their distribution across diverse ecosystems directly from genome data. By linking thallusin biosynthesis to conserved *ebo* core genes, cyclitol-derived precursor metabolism, and quinoline prenylation, this study establishes the first biosynthetic framework for a bacterial morphogen that controls macroalgal development at picomolar concentrations. Screening of 124,295 prokaryotic genomes revealed *ebo* core genes across eleven bacterial and two archaeal phyla from marine, terrestrial, and freshwater habitats, demonstrating that the capacity for thallusin biosynthesis is far more widespread than previously recognized. The discovery of thallusin production in cyanobacteria and *Actinomycetota* expands its known phylogenetic and ecological range beyond marine *Bacteroidota* and suggests that thallusin-associated signaling extends beyond algal surface microbiomes into diverse microbial communities and environmental interfaces. The previously reported observation that deletion of *ebo* genes in the cyanobacterium *N. punctiforme* impairs biosynthesis of the UV-protective pigment scytonemin (*22*) further suggests that thallusin biosynthesis or related *ebo*-associated metabolites may participate in broader cellular stress-response or regulatory processes beyond algal morphogenesis. Together, these findings reposition thallusin from a specialized marine algal morphogen to a globally distributed microbial signaling metabolite with potential functions across aquatic and terrestrial ecosystems.

## Materials and Methods

### Sampling, isolation and initial cultivation

*Ulva* sp. thalli were collected on February 7^th^, 2024, on Heligoland Island, Germany (coordinates: 54.17939, 7.88900) and were kept at 4 °C during the transfer to the lab for further processing. Artificial seawater version 2 medium (ASW_v.2_ medium), that is based on a previously published protocol (*32*), was solidified with 0.8% (w/v) Gelrite and used for the incubation of the sampled *Ulva* sp. specimens. ASW_v2_ medium contained basal artificial seawater version 2 (ASW_v.2_, 23.926 g/L NaCl, 4.008 g/L Na_2_SO_4_, 677 mg/L KCl, 98 mg/L KBr, 3 mg/L NaF, 21 mg/L Na_2_CO_3_, 168 mg/L NaHCO_3_, 10.3 g/L MgCl · 6H_2_O, 1.5 g/L CaCl_2_ · 2H_2_O, 25 mg/L SrCl_2_ · 6H_2_O), vitamins (50 µg/L *p*-aminobenzoic acid, 20 µg/L biotin, 100 µg/L pyridoxine hydrochloride, 50 µg/L thiamine hydrochloride, 50 µg/L Ca-pantothenate, 20 µg/L folic acid, 50 µg/L riboflavin, 50 µg/L nicotinamide and 20 µg/L vitamin B_12_), L1 trace elements (4.36 mg/L Na_2_EDTA · 2H_2_O, 3.15 mg/L FeCl_3_ · 6H_2_O, 178.1 µg/L MnCl_2_ · 4H_2_O; 23 µg/L ZnSO_4_ · 7H_2_O, 11.9 µg/L CoCl_2_ · 6H_2_O, 2.5 µg/L CuSO_4_ · 5H_2_O, 19.9 µg/L Na_2_MoO_4_ · 2H_2_O, 2.7 µg/L H_2_SeO_3_, 2.6 µg/L NiSO_4_ · 6H_2_O, 1.8 µg/L Na_3_VO_4_, 1.9 µg/L K_2_CrO_4_), L1 nutrients (75 mg/L NaNO_3_, 5 mg/L NaH_2_PO_4_ · H_2_O), 5 mM NH_4_Cl and 5.5 mM *N*-acetyl glucosamine as the sole carbon source. Additionally, 50 mg/L cycloheximide, 30 mg/L carbenicillin, 100 mg/L streptomycin, 200 mg/L ampicillin and 50 mg/L kanamycin were added to the medium to prevent growth of fungi and fast-growing bacteria. The pH was adjusted to 8.0 with HCl. Collected *Ulva* sp. thalli were washed twice with sterile ASW v2 medium, placed in the center of ASW v2 Gelrite plates and incubated at 18 °C. After two months of incubation, single colonies were picked from the plates and subsequently cultivated in marine M1 medium (synonym: M1H NAG ASW medium) at pH 7.5 (*12*).

### Genome sequencing and assembly, genome-based analyses and accession numbers

For the isolates UH1 and UH5 genomic DNA was extracted with the Wizard HMW DNA Extraction Kit (Promega) following the manufacturer’s instructions, but instead of 100 µL rehydration buffer, 50 µL TE buffer (10 mmol/L Tris-HCl, pH 8.0, 1 mmol/L EDTA-Na_2_, Promega) were used. DNA quantification and quality control were conducted as previously reported (*33*). For the isolate UH7 whole genome amplification (wga) was performed from 1 µL of an early exponential culture using the REPLI-g Single Cell Kit (Qiagen). The amplification reaction was performed in quarter reactions and the obtained wgaDNA was processed as previously reported (*34*). During the first DNA purification step, all volumes were downscaled accordingly. For all strains *de novo* genome assembly was performed using long-read data from Oxford Nanopore sequencing, with subsequent polishing utilizing short-read data from Illumina sequencing. For this, the sequencing reads were uploaded to the Galaxy web platform and the server available under the public domain usegalaxy.eu (*35*). Customized workflows were used for the processing of the data as previously reported (*33*). Details on the sequencing chemistry and bioinformatic workflow as well as tools, tool versions and optional parameters are stated in Table S4. Illumina sequencing was performed by Eurofins Genomics (Ebersberg, Germany). Genome sequencing of strain MS6 was performed by GATC Biotech (Konstanz, Germany) using the PacBio single-molecule real-time (SMRT) sequencing technology on the Pacific Biosciences RS sequencer (Menlo Park, CA, USA). The service included library preparation, quality control, raw read filtering, genome assembly and gene annotation. Genomic DNA was prepared from 30 mL of a late log-phase cell culture using the CTAB protocol provided by the Joint Genome Institute (JGI, Berkeley, USA).

The genome completeness was evaluated using BUSCO v. 5.8.0 (*36*) and CheckM v. 1.2.3 (*37*). Coding density and DNA G+C content were determined using CheckM v. 1.2.3. Following an initial annotation with Prokka v. 1.14.6 (*38*), the chromosome was re-oriented to the start codon of the *dnaA* gene encoding the replication initiator protein, and subjected to final re-annotation using PGAP v. 2025-05-06, build 7983 (*39*). The pangenome of selected strains was generated with anvi’o v.9 with the following parameters: minimum minbit value of 0.5, MCL (Markov Clustering) inflation of 2 and minimum amino acid sequence identity of 0% (*40*). Biosynthetic gene clusters were predicted with antiSMASH v.8.0 in relaxed mode and with all extra features activated (*41*). The annotation with precomputed orthologous groups and phylogenies was performed with eggnog-mapper v. 2.1.13 (*42*). Genome sequences have been deposited at NCBI under the following accession numbers: JBWWRM000000000 (*Maribacter* sp. UH1), JBWWRN000000000 (*Rhodopirellula* sp. UH5), JBWWRO000000000 (*Aurantibacter* sp. UH7) and JBWWRP000000000 (*Maribacter* sp. MS6).

### Cultivation procedures

*Escherichia coli* DH5α or TOP10 were used for cloning and were routinely cultivated in LB medium (5 g/L yeast extract, 10 g/L tryptone, 5 g/L sodium chloride) at 37 °C. *E. coli* BL21(DE3) was used as a host for the heterologous expression of *eboC* genes and was cultivated in TB medium. TB medium contained per liter: 12 g tryptone, 24 g yeast extract, 0.4% (v/v) glycerol, 17 mmol/L KH_2_PO_4_, 72 mmol/L K_2_HPO_4_. Antibiotics (34 mg/L chloramphenicol, 60 mg/L kanamycin or 100 mg/L ampicillin) were used for the selection of recombinant strains and maintenance of plasmids. *Stieleria maiorica* Mal15 and *Saccharomonospora* sp. CNQ-490 were cultivated in 50 mL marine M3 medium (synonym: M3H NAG ASW) (*12*) using 500 mL baffled flasks with shaking (130 rpm) at 28 °C. The medium was supplemented with 50 mg/L chloramphenicol for the cultivation of *S. maiorica* deletion mutants. Solidified M3H NAG ASW medium was prepared with 15 g/L agar that was washed three times with ddH_2_O, autoclaved separately and added shortly before pouring the plates.

### Plasmid construction

PCR-based gene amplification and plasmid construction were performed using the manufacturer’s protocols. Strains and plasmids are listed in Table S5 and oligonucleotides in Table S6. As a template, either 1 µL of a liquid culture or gDNA extracted using the DNeasy Blood & Tissue Kit (Qiagen) was used. DreamTaq PCR Mastermix Green (Thermo Scientific) was used for all check-PCR reactions and Q5 polymerase Master Mix (NEB) or S7 Fusion High-Fidelity DNA Polymerase (Biozym) was used for the amplification of sequences to be inserted in plasmids. Plasmids were constructed using restriction and ligation with FastDigest restriction enzymes and the Rapid DNA ligation kit (Thermo Scientific). All generated plasmids were verified by PCR using check primers that bind outside of the inserted sequences and by DNA sequencing at Macrogen Europe. EboC-encoding gene sequences optimized to the codon usage of *E. coli* were obtained from BioCAT GmbH already subcloned in the vector pET28a and were directly used for the transformation of *E. coli* BL21(DE3) for heterologous expression.

### Construction of *S. maiorica* gene deletion mutants

Gene deletion mutants of *S. maiorica* were constructed based on a previously published homologous recombination protocol (*43*). Briefly, 1500 bp homology regions flanking the open reading frame of the inactivation target were PCR-amplified and inserted in a plasmid up- and downstream of a resistance cassette bearing a chloramphenicol acetyltransferase (*cat*) gene. Subsequently, 2 µg of linearized plasmid DNA were used for the electroporation of competent *S. maiorica* cells. Preparation of electrocompetent cells, transformation, and cultivation of recombinant strains were performed as previously described (*43*). Due to strong strain aggregation, recombinant clones were not always immediately free of residual wild-type cells. Hence, recombinant strains were cultivated for 2-5 additional rounds in 10 mL liquid marine M1 medium supplemented with 50 mg/L chloramphenicol until a check-PCR with primers binding within the open reading frame of the deleted gene was negative.

### Morphogenetic bioassays with *Ulva* sp

#### Algal material and preparation of axenic cultures

Mature thalli of *Ulva compressa* (formerly *Ulva mutabilis* strain FSU-UM5-1) were cultured under a 17 h light / 7 h dark regime at 18 °C in *Ulva* Culture Medium (UCM). Gametogenesis was induced by mincing the thallus through a chopper and washing the thallus fragments according to standard procedures (*44*). Upon three days of induction, the released gametes were discharged, collected, and subjected to phototactic purification to separate them from bacteria. Gametes were allowed to swim toward a light source within a glass capillary under strictly sterile conditions (*45*). The final suspension was checked by PCR using universal bacterial 16S rRNA primers to confirm axenicity. The axenic cultures were inoculated with *Roseovarius* sp. MS2 and the putative thallusin producers or supernatant.

#### Preparation of bacterial cultures and test samples

Members of *Bacteroidota* were grown in HaHa_100 medium at 18 °C, while members of *Planctomycetota, Pseudomonadota* and *Actinomycetota* as well as the deletion mutants of *S. maiorica* were cultivated in marine M1 or M3 medium at 28 °C. Cells were harvested during the exponential phase and washed twice with sterile UCM.

#### Ulva bioassay for morphogenesis and thallusin activity

The *Ulva* bioassay was performed in sterile 50 mL cell culture flasks with filter caps (T25, Sarstedt AG, Germany) containing 20 mL *Ulva* culture medium (UCM) and approximately 500 purified gametes. Gametes were inoculated with washed bacterial strains (OD_620_ = 0.001), including wild-type strains and *ebo* deletion mutants, in combination with *Roseovarius* sp. MS2 for assessment of thallusin activity within the tripartite community (Fig. S1). Cultures were incubated under standard *Ulva* cultivation conditions, and morphological development was monitored microscopically over 14 days. Morphogenetic responses were classified into four categories (Table S1). Positive controls consisted of co-cultures with *Roseovarius* sp. MS2 (cytokinin-like activity) and *Maribacter* sp. MS6 (thallusin producer) to induce complete morphogenesis of *Ulva* (*46*). Additional controls included axenic cultures and cultures inoculated with either *Roseovarius* sp. MS2 or *Maribacter* sp. MS6 alone (Fig. S1).

For testing cell-free supernatants from cyanobacterial cultures, *Nostoc* sp. PCC 7120 and *Nostoc punctiforme* PCC 71302 were pre-grown in BG-11_0_ and *Synechocystis* sp. PCC 6803 in BG-11 medium (*47*) supplemented with 5 mM HEPES pH 7.5 at 30 °C with continuous illumination. The culture medium was sterile-filtered through a 0.2 µm PES membrane (Filtropur S, Sarstedt, Nümbrecht, Germany). *Ulva* cultures (20 mL) were inoculated with 100 µL of sterile-filtered supernatant for bioassay analysis. The same controls were applied as for bioassays with heterotrophic bacteria.

### Extraction, derivatization and LC-HRMS/MS analysis of thallusin

After bacterial cultivation, cultures were centrifuged to remove biomass, and the supernatant was collected and filtered through a 0.22 µm PES membrane to remove residual particles and ensure sterility. Following the addition of an internal standard, a monomethylated (−)-thallusin derivative with a shifted double bond and one reduced carboxylic acid (*10*), the cell-free supernatant was loaded onto preconditioned C_18_ solid-phase extraction cartridges (Sep-Pak Plus, Waters, UK) for metabolite enrichment. The solid-phase extraction (SPE) cartridges were washed with 10 mL MicroPure water and 4 mL of 25% (v/v) methanol, and thallusin was subsequently eluted with 4 mL of 75% methanol. Eluates were dried under a gentle stream of nitrogen and derivatized with iodomethane (MeI), resulting in complete methylation of carboxylic acid groups (*10*). After an additional C_18_-SPE cleanup step, the derivatized samples were analyzed by UHPLC-HR-MS/MS using positive electrospray ionization (ESI^+^).

### Mass spectrometric analysis of thallusin and saccharoquinoline

To investigate the presence and structural characteristics of thallusin, ultra-high-performance liquid chromatography coupled to electrospray ionization high-resolution mass spectrometry (UHPLC-ESI-HRMS) was employed. Analyses were performed on a Q Exactive™ Plus Quadrupole-Orbitrap mass spectrometer (Thermo Fisher Scientific, Dreieich, Germany) coupled to an UltiMate HPG-3400 RS binary pump. Chromatographic separation was achieved using a Kinetex C_18_ reversed-phase column (50 × 2.1 mm, 1.7 µm, 100 Å; Phenomenex, Germany) maintained at 25 °C. Ionization was carried out using a heated electrospray ionization source in positive mode. Full-scan data were acquired in total ion current (TIC) mode (*m/z* 100-1000) at a resolution of 280,000 full width at half maximum (FWHM), with an an automatic gain control (AGC) target of 3 × 10⁶, a maximum injection time of 200 ms, and an acquisition window of 4.0-8.0 min. In parallel, targeted selected ion monitoring (tSIM) was used to detect underivatized thallusin (*m/z* 458.2173 ± 0.2), derivatized thallusin trimethyl ester (m/z 500.2643 ± 0.2), monomethylated saccharoquinoline (*m/z* 440.2431 ± 0.2), and the methylated internal standard (*m/z* 472.2694 ± 0.2). tSIM data were acquired at 70,000 FWHM, with an AGC target of 2 × 10⁵ and a maximum injection time of 200 ms. Source parameters were set as follows: sheath gas 60, auxiliary gas 20, sweep gas 5, spray voltage 3.0 kV, capillary temperature 360 °C, S-lens RF level 50, and auxiliary gas heater temperature 400 °C. For structural characterization, MS/MS fragmentation of derivatized thallusin ([M+H]⁺, m/z 500.2643 ± 0.2) and isotopologues were performed using higher-energy collisional dissociation at a normalized collision energy of 32. Fragment ions were detected in Orbitrap mode at 35,000 FWHM, with an AGC target of 2 × 10⁴ and a maximum injection time of 100 ms. Data were processed using Xcalibur software (Thermo Fisher Scientific). Chromatographic separation was employed with eluent A (2% acetonitrile, 0.1% formic acid in water) and eluent B (acetonitrile with 0.1% formic acid). The gradient was programmed as follows: 0.2 min, 0% B (0.4 mL/min); linear increase to 100% B by 8.0 min (0.675 mL/min); 8.0-12.0 min, 100% B (0.675 mL/min); 12.0-12.3 min, return to 0% B (0.4 mL/min); and 12.3-15.0 min, equilibration at 0% B (0.4 mL/min). Samples (3 µL) were injected using a WPS-3000 autosampler at 10 °C. The trimethylated analyte was detected at *m/z* 500.2643 [M+H]^+^ with a retention time of 6.48 min and was compared with synthetic standards of thallusin. The synthetic internal standard showed a corresponding methylated ion at *m/z* 472.2694, confirming quantitative methylation. MS/MS spectra displayed diagnostic fragment ions characteristic of the drimane-type sesquiterpenoid and quinoline carboxylate moieties (*m/z* 296.0765 [C₁₃H₁₂O₇N]⁺), consistent with the thallusin structure (*8*).

### Uptake experiments with stable isotopically labeled precursors of thallusin

Uptake experiments were performed using stable isotope-labeled compounds (Table S7) to assess their incorporation into thallusin. Compounds were added to 50 mL sterile HaHa_100 medium (*48*) to a final concentration of 1.0 mmol/L and sterile-filtered through a PES membrane. *Maribacter* sp. MS6 cultures were inoculated to an initial OD_620_ of 0.01 and incubated at 20 ± 1 °C with shaking. After 3 days (OD₆₂₀ = 0.25), cultures were centrifuged (12,000 × g, 20 min, 10 °C), and the resulting supernatants were subjected to C_18_-SPE, followed by MeI derivatization as described above and analyzed by UHPLC-HR-MS/MS to detect isotopic incorporation into thallusin.

### Chemical synthesis of 6,7,8-trihydroxyquinaldic acid (6,7,8-HQA) ethyl ester

The farnesylation substrate (compound **7**) was synthesized as the ethyl ester (to enhance solubility and permeability) from commercially available 2,3,4-trihydroxybenzoic acid (compound **1**, Sigma, 98%) as shown in Fig. S10. In brief, the starting compound was initially perbenzylated (86%). The resulting ester (compound **2**) was saponified and subjected to a Hofmann degradation by using diphenylphosphoryl azide (DPPA), followed by hydrolysis of the intermediate carbamate (61%). The aniline obtained (compound **3**) was converted to quinoline (compound **6**) by using an iodine-catalyzed Doebner-Miller cyclization (*49*), with unsaturated ester (compound **5**) as a reagent that was obtained from aldehyde (compound **4**) by using a Wittig-Horner-Emons reaction (91%). Hydrogenolysis of the benzyl ethers, followed by purification on RP silica, gave 6,7,8-HQA ethyl ester (compound **7**, 96% yield, 38% overall). Experimental details including chemical compound characterization of all new compounds can be found in the Supplementary text section.

### Preparation of enzyme extracts and enzyme assay

Plasmid-borne expression of *eboC* genes in *E. coli* was performed as previously described (*19*). Cultures were then centrifuged, the cell pellet was resuspended in Tris-HCl buffer (50 mmol/L, pH 7.8) supplemented with dithiothreitol (DTT, 10 mmol/L), and sonicated on ice (12 min, 100% C, 40% A, 2 seconds on and 3 seconds off intervals) by using a Hielscher UP200St ultrasonic processor. The first cell-membrane fraction was obtained by centrifugation (12,000 g, 20 min, 4 °C), while the enriched protein fraction, likely embedded in lipidic membrane components, required ultracentrifugation of the crude protein lysates (240,000 g, 90 min). The obtained protein pellet was resuspended in the same buffer as used for the bioassays. The protein concentration was measured by using Bradford Reagent (Sigma-Aldrich, St. Louis, MO, USA). Enzyme assays were performed using a standard reaction mixture (100 µL) by adding 50 µL Tris-HCl (50 mmol/L, pH 7.8) containing MgCl_2_, farnesyl pyrophosphate, aromatic substrate (each 1 mmol/L) and 50 µL of membrane-bound protein aliquots with a total protein content of 0.4 mg. The reaction mixtures were incubated at 30 °C for 2 h and subsequently extracted three times with ethyl acetate (450 µL). The solvent was removed in vacuo, the residue was dissolved in methanol (100 µL) and analyzed by HRMS/MS. Protein denaturation was performed at 95 °C for 10 min (negative control).

### HRMS-MS analysis of the enzymatic products of the EboC assay

UHPLC-HRMS measurements were carried out on a Vanquish Flex UHPLC system (Thermo Fisher Scientific) combined with an Orbitrap Exploris 120 mass spectrometer equipped with a HESI source (Thermo Fisher Scientific). Metabolites were separated using reverse phase liquid chromatography at 40 °C using a Kinetex^®^ C18 column (50 × 2.1 mm, particle size 1.7 µm, 100 Å, Phenomenex) preceded by a C18 SecurityGuard^TM^ ULTRA guard cartridge (2.1 mm, Phenomenex). Mobile phases consisted of H_2_O + 0.1 % formic acid (A) and acetonitrile (ACN) + 0.1% formic acid (B). 5 µL sample, concentrated at 50 µg/mL, was injected into a gradient as follows: 0-1 min, 5% B; 1-10 min, 5-97% B; 10-12 min, 97% B; 12-13 min, 97-5% B; 13-15 min, 5% B at a constant flow rate of 0.3 mL/min. Data-dependent acquisition of MS^2^ spectra was performed in positive mode. HESI parameters were set to 50 AU sheath gas flow, 13 AU auxiliary gas flow, 1 AU sweep gas flow, 3.4 kV (+) spray voltage, 300 °C vaporizer temperature and 320 °C ion transfer tube temperature. MS^1^ full-scan parameters were set to data type-centroid, *m/z* 150-1,500 scan range, resolving power 60,000 at *m/z* 200, 1 micro-scan, 100 ms max. injection time, 1 × 10^6^ automated gain control, 70 % RF lens, dynamic exclusion filter-auto and isotope exclusion filter-assigned. Up to four MS^2^ spectra per MS^1^ survey scan were recorded with the following parameters: data type-centroid, scan range-auto, resolving power 30,000 at *m/z* 200, 1 micro-scan, 100 ms max. injection time, 1 × 10^5^ automated gain control, 1.2 *m/z* isolation window, collision energy type-normalized with a stepwise increase from 20 to 30 to 40%.

### Mapping the prevalence of thallusin biosynthetic genes across the prokaryotic tree of life

To analyze the potential for thallusin biosynthesis, 124,295 non-redundant genomes were downloaded from mOTUs-db (http://www.motus-db.org) (*50*). These genomes are representatives of species-level clustered operational taxonomic units (mOTUs) and were selected from a collection of 3.75M systematically processed prokaryotic genomes, including MAGs, SAGs, and isolates, obtained from 118K global samples. Each mOTUs-db genome was taxonomically classified with GTDB R220 using GTDB-Tk v. 2.4 (*51*), and protein-coding genes were predicted using Prodigal v. 2.6.3 (parameters: -c -m -g 11 -p single) (*52*). The translated genomes were used as query to search with MMseqs2 (*53*) in easy-search mode (Git commit 8ef870f95af2a3ee474c2cdbb845f5f007fe5be6; default parameters: -s 5.7, -e 1.000E-03) for homology against nine target protein sequences associated with thallusin biosynthesis (EboA-F from *Maribacter stanieri* DSM 19891 (RefSeq accession GCF_900112245.1) and Ino1-3 from *Saccharomonospora* sp. CNQ-490 (RefSeq accession GCF_000527075.1). The first objective was to describe the composition of thallusin biosynthetic gene clusters and thereby enabling the assignment of core and variable components. Thallusin clusters were defined as regions containing at least four of the nine target proteins, with adjacent MMSeqs2 hits separated by no more than six open reading frames. This information was extracted from query names and the MMSeqs2 output using a custom R (v. 4.5.3) script available on GitHub (https://github.com/martsper/thallusin-motus-analysis). For all downstream analyses, only phyla harboring at least ten *ebo* clusters were considered.

The potential for thallusin biosynthesis in bacteria at the order level was analyzed next to identify the core genes. Owing to the observation that the identified core genes *eboB*, *eboC*, *eboE* and *eboF* are scattered throughout the genomes in certain phyla, the potential for thallusin biosynthesis was deduced from the presence of all four core genes at any location in each genome. For visualization, the bacterial GTBD R220 phylogenetic tree was downloaded from the GTDB webpage (https://data.gtdb.aau.ecogenomic.org) (*51*), loaded in R and pruned to branches in which GTDB identifiers match those of the mOTUs-db genome collection. Branches were collapsed at the order level, keeping only orders that harbor at least ten genomes and in which at least one genome contains all four core genes (*eboB*, *eboC*, *eboE* and *eboF*). Operations were performed in R using the package ape (v. 5.8.1) (*54*). The percentage prevalences of genomes with all four core genes per order were plotted onto the pruned GTDB tree using iTOL (v. 7.5.1) (*55*).

## Acknowledgements

The authors thank William Fenical (Scripps Institution of Oceanography, San Diego, USA) for providing *Saccharomonospora* sp. CNQ-490 and Vicente Mariscal (CSIC, Universidad de Sevilla, Spain) for providing *Nostoc punctiforme*. The authors thank Carmen E. Wurzbacher and René Maskos (Friedrich Schiller University, Jena, Germany) for their assistance with genomic DNA isolation and genome sequencing and Maja Bocker (Friedrich Schiller University, Jena, Germany) for assistance in sample preparation for thallusin identification via LC-MS. We thank H.-J. Ruscheweyh (ETH Zurich, Switzerland) for bioinformatic support with mOTUs-db. The authors thank the Jena School for Microbial Communication (JSMC) for continuous support. We thank H.-J. Ruscheweyh (ETH Zurich, Switzerland) for bioinformatic support with mOTUs-db.

## Funding

This study was funded by the German Research Foundation (DFG, Deutsche Forschungsgemeinschaft) under Project-ID 239748522 (CRC ChemBioSys, subprojects A01, A02, A06, A07, B08) and Germany’s Excellence Strategy under Project-ID 390713860 - EXC 2051 (to CJ, HDA). The Konrad-Adenauer Foundation is acknowledged for providing a doctoral fellowship to JFU, which supported this research. ShSu acknowledges funding from the Swiss National Science Foundation (project 205320_215395).

## Author contributions

Conceptualization: N.K., J.F.U., C.J., T.W.; Investigation: N.K., J.F.U., My.St., Y.L., M.K.D., M.W., H.H., J.H., R.N.; Formal analysis: N.K., Ma.Sp.; Visualization: N.K., J.F.U., Ma.Sp., H-D.A., C.B., T.W.; Resources: J.A.Z.Z.; Supervision: Se.Sa., Sh.Su., H-D.A., C.B., C.J., T.W.; Funding acquisition: C.B., Se.Sa., H-D.A., C.B., C.J., T.W.; Project administration: C.J., T.W.; Writing - original draft: N.K., C.J., T.W.; Writing - review & editing: all authors. All authors have read the final version of the manuscript and agree with the submission.

## Conflicts of interest

The authors declare that there is no conflict of interest.

## Data, code, and materials availability

Genome sequences have been deposited at NCBI under the following accession numbers: JBWWRM000000000 (*Maribacter* sp. UH1), JBWWRN000000000 (*Rhodopirellula* sp. UH5), JBWWRO000000000 (*Aurantibacter* sp. UH7) and JBWWRP000000000 (*Maribacter* sp. MS6). The custom R script used for the identification of *ebo* genes in prokaryotic genomes is available at GitHub: https://github.com/martsper/thallusin-motus-analysis. All data needed to evaluate the conclusions in the paper are present in the paper and/or the Supplementary Materials.

## Supplementary Materials

Materials and Methods

Supplementary Text

Figs. S1 to S20

Tables S1 to S7

Supplementary References

## Notes

https://www.ncbi.nlm.nih.gov/bioproject/1443044/

https://github.com/martsper/thallusin-motus-analysis

### Competing Interest Statement

The authors have declared no competing interest.

