## Supplementary material for "Conserved bacterial genes for biosynthesis of the algal morphogen thallusin span land and sea": Fig. S

### Supplementary Text

**Preparation of benzyl 2,3,4-tris(benzyloxy)benzoate (Compound 2, Fig. S10)**

2,3,4-Trihydroxybenzoic acid (2.00 g, 11.8 mmol, 1.0 equiv.) and $K_2CO_3$ (16.3 g, 118 mmol, 10.0 equiv.) were placed in a 250 mL round bottom flask, and DMF (25 mL) was added. The suspension was cooled to 0 °C with stirring, and BnBr (13.98 mL, 117.6 mmol, 10.0 equiv.) was added dropwise. The mixture was then heated to 80 °C with stirring for 16 h. After cooling to room temperature, the mixture was diluted with cold water (100 mL). The resulting precipitate was collected by filtration and dried under vacuum. The crude product was purified by flash column chromatography on silica gel (100 g silica gel, 20% to 40% EtOAc in petroleum ether), to afford 5.36 g of the title compound as a colorless solid (81% yield). mp 63 °C. $R_f$ = 0.51 (15% EtOAc in hexane). **$^1$H NMR** (300 MHz, $CDCl_3$) δ 7.57 (d, *J* = 8.8 Hz, 1H), 7.36 - 7.16 (m, 20H), 6.70 (d, *J* = 8.8 Hz, 1H), 5.23 (s, 2H), 5.06 (s, 2H), 5.01 (s, 2H), 4.94 (s, 2H) (Fig. S11); **$^{13}$C NMR** (75 MHz, $CDCl_3$) δ 165.4, 156.7, 153.9, 142.6, 137.3, 136.2, 136.2, 128.8, 128.7, 128.7, 128.6, 128.4, 128.3, 128.2, 128.2, 128.1, 128.0, 127.5, 127.3, 118.8, 108.8, 76.3, 75.6, 70.9, 66.7 (Fig. S12); **IR:** $\tilde{\nu}$ max 2975, 2855, 1714, 1584, 1487, 1440, 1239, 1110, 955, 787 cm$^{-1}$; HRMS (ESI) m/z 531.2154 [M + H]$^+$; calculated for $[C_{35}H_{30}O_5 + H]^+$: 531.2166.

**Procedure for the preparation of 2,3,4-tris(Benzyloxy)aniline (Compound 3, Fig. S10)**

To a solution of ester **2** (1.0 g, 1.88 mmol, 1.0 equiv.) in a mixture of THF and $H_2O$ (1:1, 20 mL), was added potassium carbonate (237 mg, 5.65 mmol, 3.0 equiv.). The mixture was stirred at 70 ºC for 12 h. After completion of the reaction (monitored by TLC), the solvent was evaporated under reduced pressure. The residue was partitioned between ethyl acetate (100 mL), water (50 mL), and 1N HCl (5 mL). The organic extract was washed with brine (20 mL), dried with $Na_2SO_4$, and concentrated under reduced pressure to afford the crude carboxylic acid.

The crude residue was re-dissolved in 1,4-dioxane (15 mL). Then diphenyl-phospyhoryl azide (DPPA, 486 $\mu$L, 2.26 mmol, 1.2 equiv.) and $Et_3N$ (394 $\mu$L, 2.83 mmol, 1.5 equiv.) were added dropwise at room temperature. The resulting mixture was stirred at room temperature for 30 minutes. Subsequently, EtOH (7.5 mL) was added, and the mixture was stirred for 2 hours. After completion of the reaction (monitored by TLC), the solvent was evaporated under reduced pressure. A mixture of EtOH and $H_2O$ (10:1, 10 mL) was added, followed by KOH (528 mg, 9.42 mmol, 5.0 equiv.), and the mixture was heated at 110 °C for 2 hours. The solvent was evaporated under reduced pressure, and the residue was redissolved in $CH_2Cl_2$ (50 mL). The solution was washed with 5% citric acid (10 mL), 5% $NaHCO_3$ solution (10 mL), and brine (10 mL), then dried with $Na_2SO_4$ and concentrated under reduced pressure. The crude product was purified by flash column chromatography (80 g silica gel, 10% to 30% EtOAc in petroleum ether) to afford 473 mg of the title aniline **3** as a pale-brown gel (61%). $R_f$ = 0.36 (30% EtOAc in hexane). **$^1$H NMR** (300 MHz, $CDCl_3$) δ 7.59 - 7.41 (m, 15H), 6.72 (d, *J* = 8.7 Hz, 1H), 6.49 (d, *J* = 8.7 Hz, 1H), 5.23 (s, 2H), 5.17 (s, 2H), 5.13 (s, 2H) (Fig. S13); **$^{13}$C NMR** (75 MHz, $CDCl_3$) δ 145.4, 143.3, 140.8, 137.9, 137.8, 137.7, 135.5, 128.7, 128.6, 128.6, 128.6, 128.4, 128.3, 128.1, 127.9, 127.7, 111.6, 110.1, 75.6, 75.1, 72.4 (Fig. S14); **IR:** $\tilde{\nu}$ max 3464, 3387, 2960, 1529, 1325, 1226, 986, 742 $cm^{-1}$; HRMS (ESI) m/z 412.1916 $[M + H]^+$; calculated for $[C_{27}H_{25}NO_3 + H]^+$: 412.1907.

**Procedure for the preparation of Ethyl (*E*)-4,4-dimethoxybut-2-enoate (Compound 5, Fig. S10)**

To a solution of 60% (w/w) 2,2-dimethoxyacetaldehyde (3.10 mL, 17.8 mmol, 1.0 equiv.) in THF was added triethylphosphonoacetate **4** (4.00 g, 17.8 mmol, 1.0 equiv.) and $K_2CO_3$ (2.96 g, 21.4 mmol, 1.2 equiv.). The mixture was stirred at room temperature for 5 min. Water (5.4 mL) was added, and stirring was continued at room temperature for 16 h. After completion of the reaction, brine (60 mL) was added. The mixture was extracted with $Et_2O$ (100 mL). The organic extract was dried with $Na_2SO_4$ and concentrated under reduced pressure to obtain 2.83 mL of the title ester **5** as a colorless oil (91% yield), that was used without further purification. $R_f$ = 0.48 (10% EtOAc in hexane). **$^1$H NMR** (300 MHz, $CDCl_3$) δ 6.78 (dd, *J* = 15.9, 4.0 Hz, 1H), 6.15 (dd, *J* = 15.9, 1.4 Hz, 1H), 4.96 (dd, *J* = 4.0, 1.4 Hz, 1H), 4.23 (q, *J* = 7.2 Hz, 2H), 3.35 (s,

6H), 1.31 (t, $J$ = 7.1 Hz, 3H) (Fig. S15); **$^{13}$C NMR** (75 MHz, $CDCl_3$) δ 165.9, 142.5, 124.7, 100.5, 67.9, 60.6, 52.8, 25.6, 14.1 (Fig. S16); **IR:** $\tilde{\nu}$ max 2981, 2910, 2844, 1719, 1664, 1349, 1267, 1162, 1041, 982, 857 $cm^{-1}$.

**Synthesis of Ethyl 6,7,8-tris(benzyloxy)quinoline-2-carboxylate (Compound 6, Fig. S10)**

To a solution of aniline **3** (200 mg, 0.486 mmol, 1.0 equiv.) in DMSO (6 mL) was added a stock solution of TFA in DMSO (1.0 M, 10 µL, 10 mol%), followed by a stock solution of iodine in DMSO (0.1 M, 10 $\mu$L, 1 mol%). The resulting mixture was heated to 80 °C. Then a solution of ethyl (2*E*)-4,4-dimethoxy-2-butenoate (**5**, 103 mg, 0.51 mmol, 1.05 equiv.) in DMSO (3 mL) was added slowly over 2 hours by using a syringe pump. The resulting mixture was stirred at 80°C for 3 more hours. After completion of the reaction, $H_2O$ (5 mL) was added and the mixture was extracted with EtOAc (3 x 20 mL). The combined organic extracts were washed with water (2 x 10 mL), brine (2 x 10 mL), dried with $Na_2SO_4$, and concentrated under reduced pressure. The residue was purified by flash column chromatography (20 g silica gel, 10% to 25% EtOAc in petroleum ether), to afford 192 mg of the title compound as a yellow resin (76% yield). $R_f$ = 0.41 (30% EtOAc in hexane). **$^{1}$H NMR** (300 MHz, $CDCl_3$) δ 8.13 (d, $J$ = 0.9 Hz, 2H), 7.78 - 7.75 (m, 2H), 7.51 - 7.31 (m, 13H), 7.02 (s, 1H), 5.51 (s, 2H), 5.24 (d, $J$ = 5.0 Hz, 4H), 4.56 (q, $J$ = 7.2 Hz, 2H), 1.56 (t, $J$ = 7.1 Hz, 3H) (Fig. S17); **$^{13}$C NMR** (75 MHz, $CDCl_3$) δ 165.9, 154.3, 148.4, 145.1, 144.5, 139.4, 137.8, 137.4, 136.1, 135.4, 129.0, 128.7, 128.3, 128.3, 128.3, 128.1, 128.0, 127.5, 127.4, 120.7, 102.2, 76.8, 76.0, 70.9, 61.8, 14.4 (Fig. S18); **IR:** $\tilde{\nu}$max 3063, 2989, 2875, 1721, 1604, 1460, 1134, 1021, 857, 774, 709 $cm^{-1}$; HRMS (ESI) m/z 542.1914 $[M + Na]^+$; calculated for $[C_{33}H_{29}NO_5 + Na]^+$: 542.1938.

**Procedure for the preparation of Ethyl 6,7,8-trihydroxyquinoline-2-carboxylate (Compound 7, Fig. S10)**

In an oven-dried round-bottom flask, the tribenzylated quinolone **6** (50 mg, 0.096 mmol; 1.0 equiv) was dissolved in ethanol (2 mL) and EtOAc (2 mL) under $N_2$ atmosphere. Pd/C (50 mg, 10 mol% (w/w)) was added in portions, and the suspension was stirred for 10 min at room temperature. Then, the $N_2$ atmosphere was exchanged for $H_2$ by purging, and the reaction mixture was stirred for 3 h under $H_2$ atmosphere (1 bar). Upon completion of the reaction (TLC monitoring), the mixture was filtered and concentrated. The crude product was washed with pentane and then purified by using a Bakerbond™ spe Octadecyl column (C18, Part no: 7020, 03), eluting with a water/acetonitrile gradient, to obtain 23 mg of the title compound **7** as a yellow gel (96% yield). $R_f$ = 0.11 (75% EtOAc in hexane). **$^1$H NMR** (300 MHz, DMSO-$D_6$) δ 9.13 (brs, 3H), 8.15 (d, *J* = 8.5 Hz, 1H), 7.83 (d, *J* = 8.5 Hz, 1H), 6.81 (s, 1H), 4.39 (q, *J* = 7.1 Hz, 2H), 1.37 (t, *J* = 7.0 Hz, 3H) (Fig. S19); **$^{13}$C NMR** (75 MHz, DMSO-$D_6$) δ 165.6, 151.0, 143.0, 139.6, 136.1, 135.1, 135.0, 124.7, 119.1, 99.8, 61.5, 14.7 (Fig. S20); **IR:** $\tilde{\nu}$max 3640, 3194, 2985, 2899, 1723, 1645, 1599, 1260, 1139, 1080, 951, 854, 747 $cm^{-1}$; HRMS (ESI) m/z 272.0514 $[M + Na]^+$; calculated for $[C_{12}H_{11}NO_5 + Na]^+$: 272.0529.

### Supplementary Figures

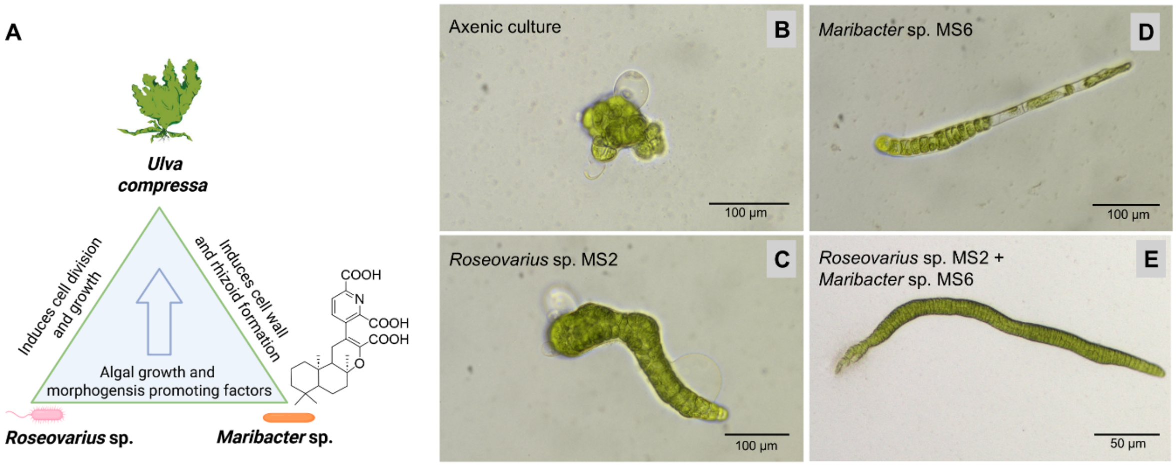


**Fig. S1. *Ulva* morphogenetic bioassay.** Biotest with *Ulva compressa* and resulting morphotypes depending on the presence of bacterial producers of a yet uncharacterized compound with cytokinin-like activity (produced by *Roseovarius* sp. MS2) or of thallusin (e.g., produced by *Maribacter* sp. MS6). **(A)** Schematic representation of the tripartite community formed by the green macroalga *Ulva compressa* and its associated bacteria *Roseovarius* sp. and *Maribacter* sp. Right panel: Light micrographs illustrating characteristic morphotypes (phenocopies) of the slender-type mutant of *U. compressa* under different bacterial conditions. **(B)** Axenic culture without bacterial supplementation, displaying callus-like thallus growth with colorless cell wall protrusions. **(C)** Culture with *Roseovarius* sp. only, characterized by pronounced cell wall protrusions. **(D)** Culture with *Maribacter* sp. only, showing altered cellular architecture and signs of cell degradation. **(E)** Co-cultivation with *Maribacter* sp. and *Roseovarius* sp., restoring complete thallus morphogenesis (see also Table S1 for details).

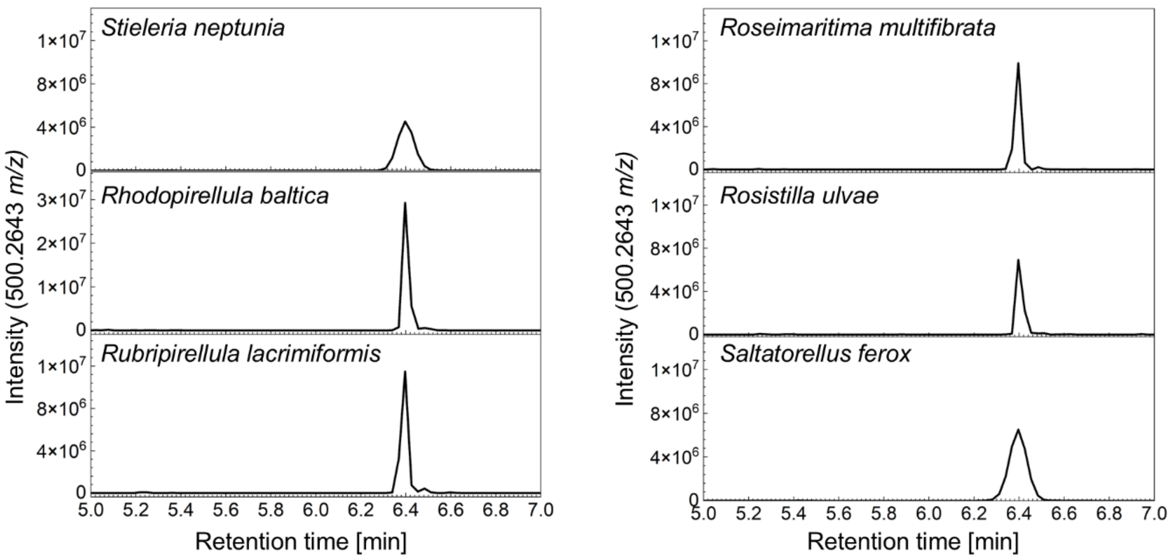


**Fig. S2. Thallusin production in different planctomycetal strains.** Extracted ion chromatograms (EIC; *m/z* 500.2643) of derivatized thallusin in selected strains belonging to the phylum *Planctomycetota*. The tested strains, including *Rhodopirellula baltica*, *Rubripirellula lacrimiformis*, *Roseimaritima multifibrata*, *Rosistilla ulvae*, and *Saltatorellus ferox*, show a characteristic peak at ~6.4 min, indicating thallusin production. These results demonstrate that thallusin biosynthesis is widespread among phylogenetically diverse planctomycetes.

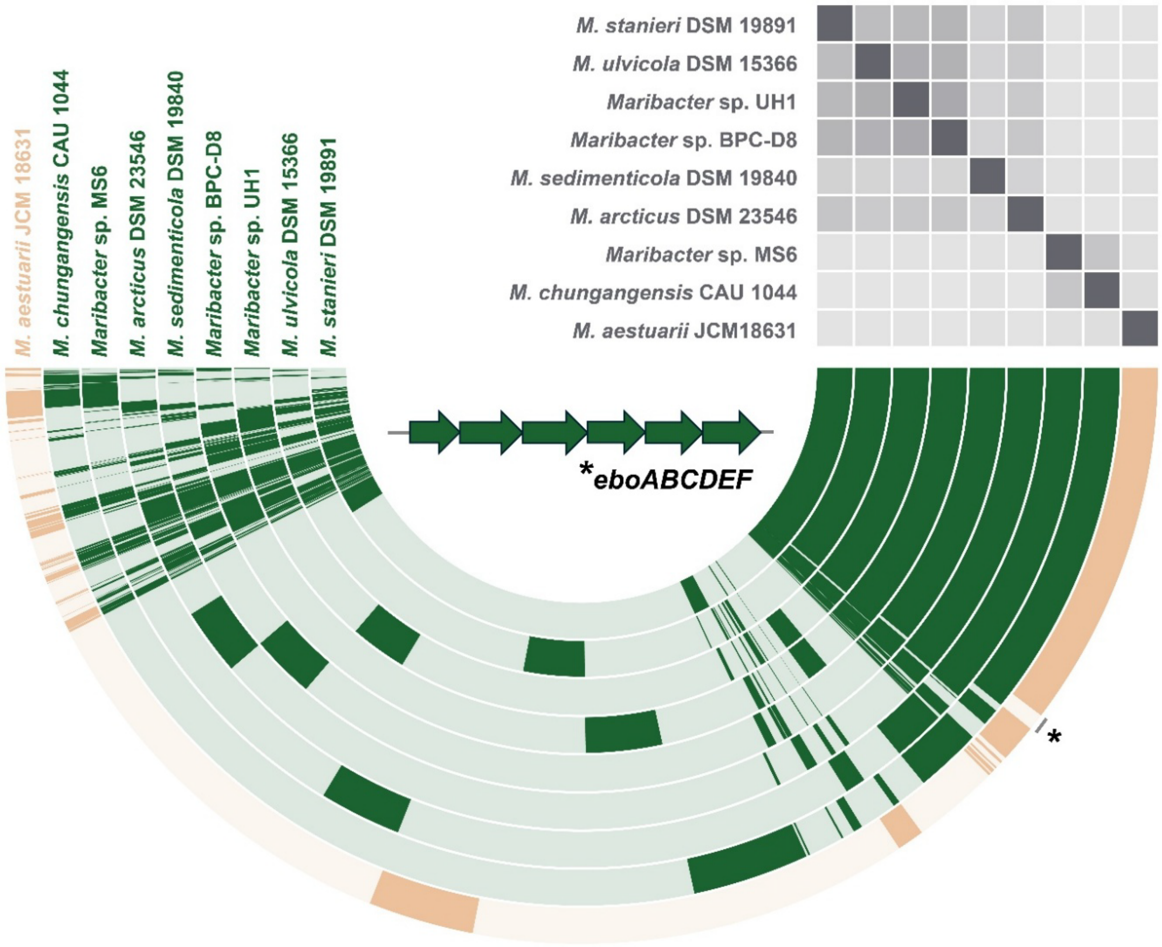


**Fig. S3. Pangenome of thallusin-producing and non-producing *Maribacter* spp.** Pangenome of eight thallusin-producing (green) and one non-producing (beige) *Maribacter* strains. Each open circle represents the pangenome of all strains but is colored darker when the gene is present in the respective genome. The heatmap in the upper right corner shows the relationship of the strains based on average nucleotide identity (ANI) values (light gray = 70% to dark gray = 100%). The asterisk indicates the gene clusters exclusively present in thallusin producers but absent in the non-producer, among which the *ebo* operon was detected. NCBI RefSeq accession numbers of the analyzed genomes are provided in Table S2. The entire set of hits and the putative COG24 annotation is provided in Table S3.

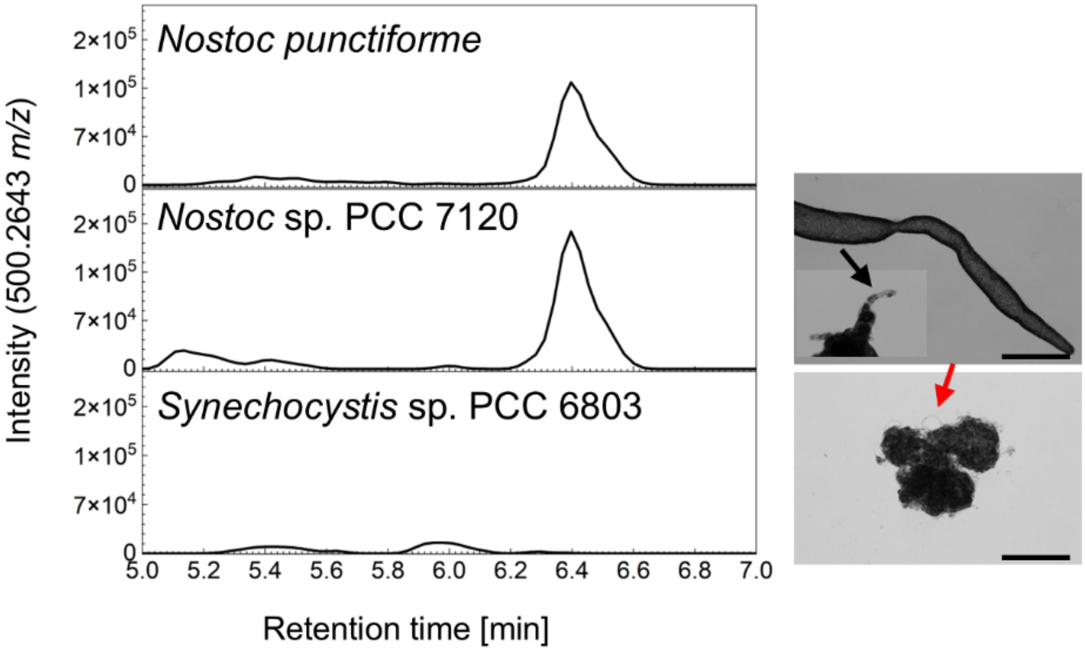


**Fig. S4. Detection of thallusin in cyanobacterial supernatants.** Extracted ion chromatograms (EIC; *m/z* 500.2643) of derivatized thallusin from sterile-filtered culture supernatants of selected cyanobacteria. *Nostoc punctiforme* PCC 73102 and *Nostoc* sp. PCC 7120 show a characteristic peak at ~6.4 min, indicating secretion of thallusin into the medium, whereas *Synechocystis* sp. PCC 6803 lacks a corresponding signal. Thallusin production correlates with the presence of *ebo* genes in producer strains, supporting a conserved role of this gene set in thallusin biosynthesis across phylogenetically distant lineages. Photographs of the *Ulva* morphogenetic bioassay in the presence of cyanobacterial supernatants and *Roseovarius* sp. MS2 (scale bar = 100 µm) are shown. Supernatants of *Nostoc* restore normal morphogenesis of *Ulva compressa*, including cell wall differentiation and rhizoid formation (inset, black arrow), consistent with thallusin activity, whereas *Synechocystis* supernatants fail to complement morphogenesis, and cell wall protrusions are visible (red arrow).

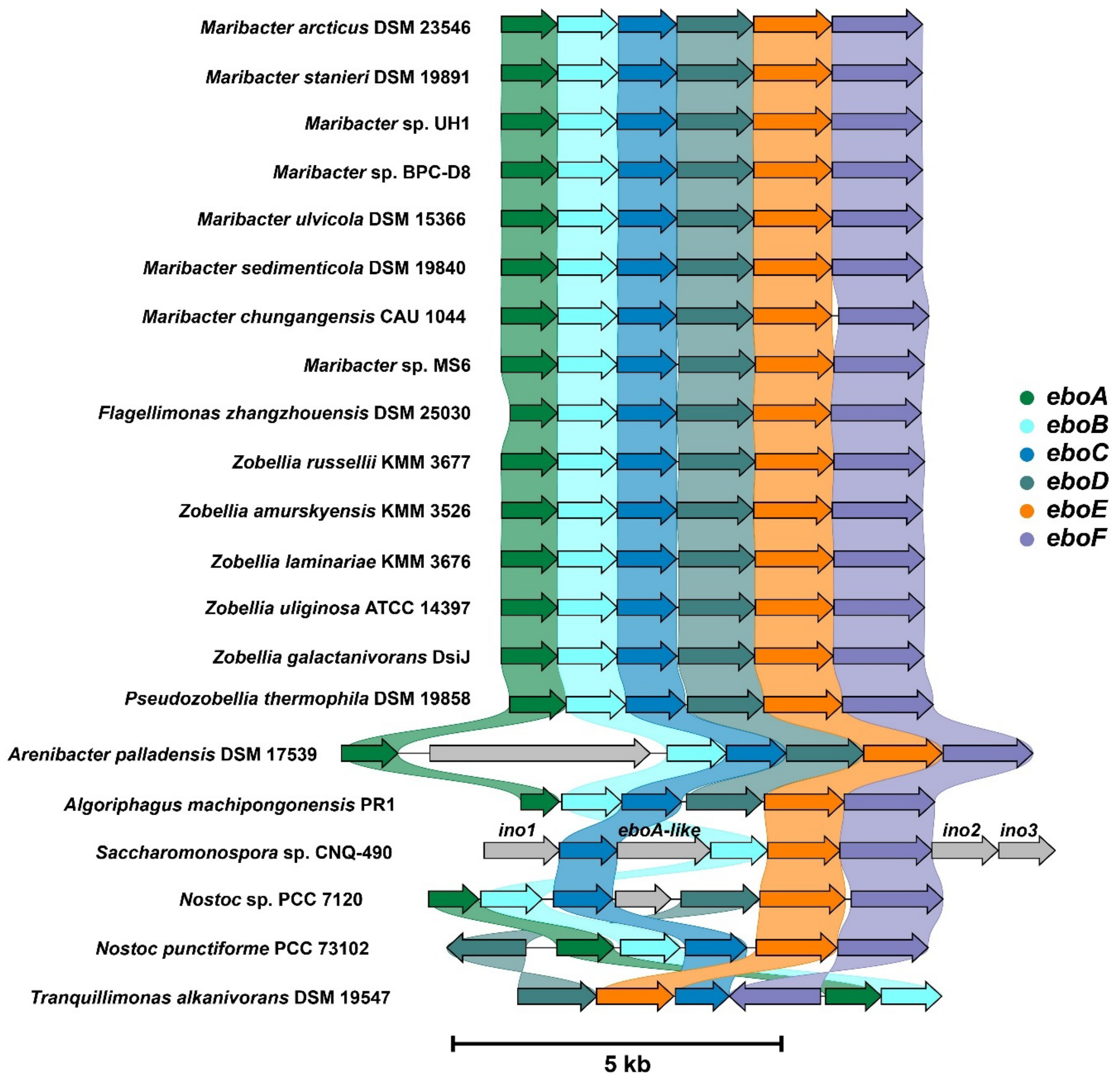


**Fig. S5. Comparison of the *ebo* gene clusters in confirmed thallusin producers.** Arrangement of *ebo* genes in analyzed genomes of thallusin-producing members of the phyla *Bacteroidota* (*Maribacter*, *Flagellimonas*, *Zobellia*, *Pseudozobellia*, *Arenibacter*, *Algoriphagus*), *Actinomycetota* (*Saccharomonospora*), *Cyanobacteriota* (*Nostoc*) and *Pseudomonadota* (*Tranquillimonas*). Due to the scattered distribution of the genes in the phylum *Planctomycetota*, the identified producer strains were not included in the analysis.

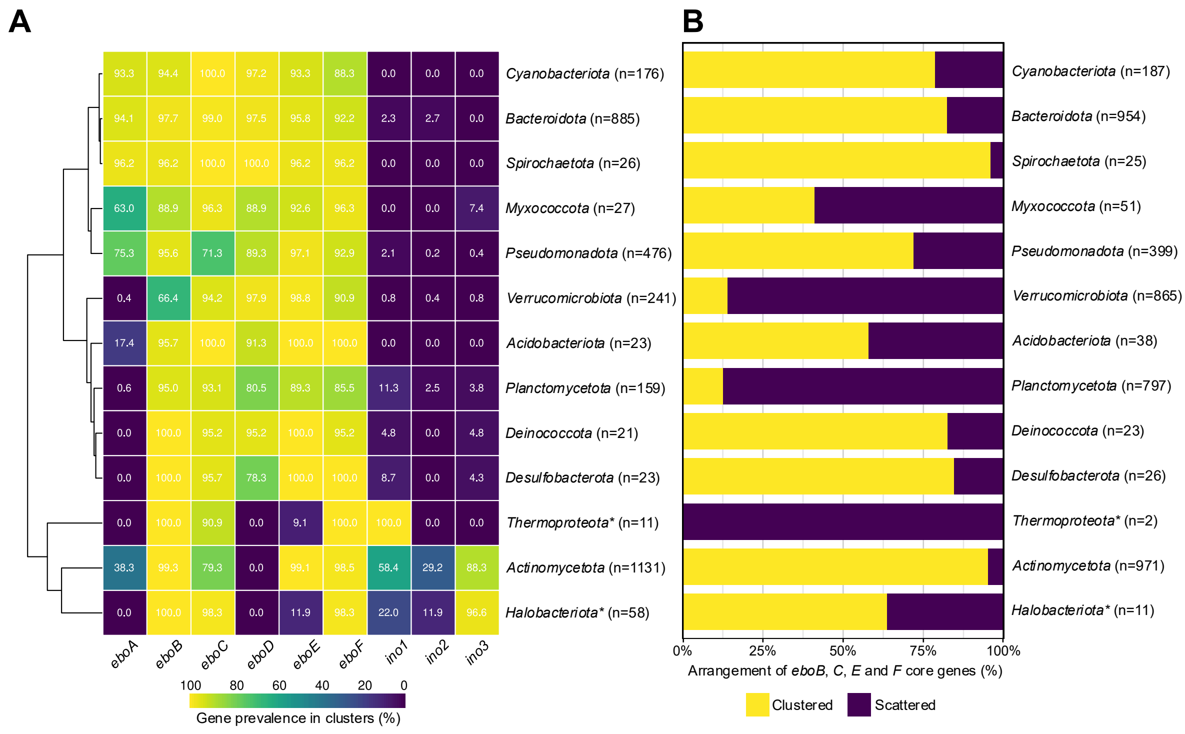


**Fig. S6. Cluster analysis of 124,295 mOTUs-db genomes identified thallusin biosynthesis gene content across 11 bacterial and two archaeal phyla.** (**A**) Identification of thallusin biosynthesis core genes (*n* = number of genomes with at least one *ebo* cluster). (**B**) Co-localization of the core genes within clusters (*n* = number of genomes containing all four core genes). In total, 4,405 of 124,295 genomes contained all four core genes (*eboB*, *eboC*, *eboE* and *eboF*) at any genomic location, of which 2,513 had the four core genes co-localized within a cluster (gap ≤ 6 ORFs between adjacent hits). Because MAGs may be fragmented, the number of genomes with co-localized core genes is likely underestimated.

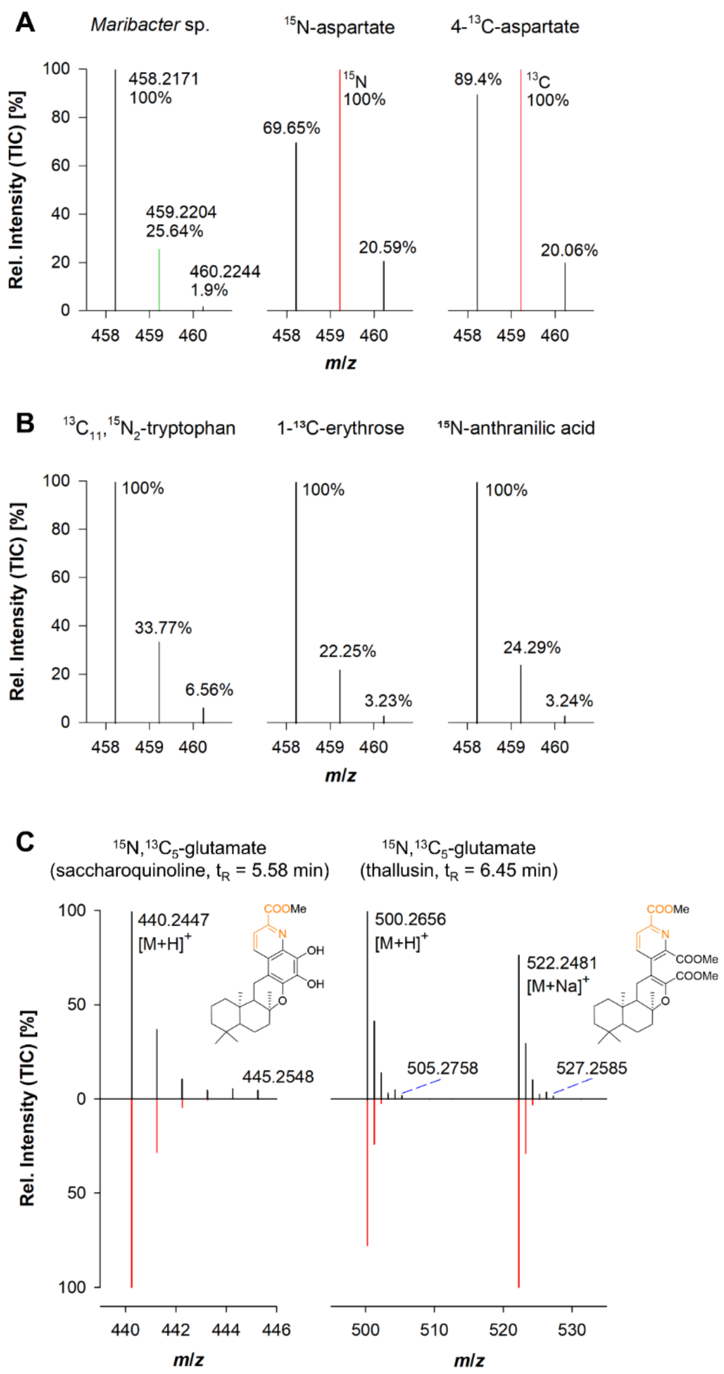


**Fig. S7. Isotope labeling with potential precursors of thallusin reveals indirect incorporation into thallusin.** (**A**) Mass spectra of thallusin (*m/z* 458.2171) in *Maribacter* sp. MS6 cultures supplemented with labeled precursors. Feeding with $^{15}N$-aspartate and $^{13}C_4$-aspartate results in strong isotopic enrichment (red line), confirming efficient incorporation into thallusin. (**B**) In contrast, labeling with alternative precursors ($^{13}C_{11}$,$^{15}N_2$-tryptophan, $^{13}C$-erythrose, $^{15}N$-anthranilic acid) results in only minor isotopic shifts, indicating limited or indirect contribution to thallusin biosynthesis. (**C**) Feeding with $^{15}N$,$^{13}C_5$-glutamate leads to detectable but partial isotope incorporation into both the monomethylated derivative of saccharoquinoline (*m/z* 440.2447) and the fully methylated thallusin (*m/z* 500.2656 $[M+H]^+$, 522.2481 $[M+Na]^+$), demonstrating that glutamate contributes indirectly to thallusin biosynthesis, likely via conversion into central metabolic intermediates such as aspartate (compare with Fig. 3). The depicted structure corresponds to the neutral molecule [M].

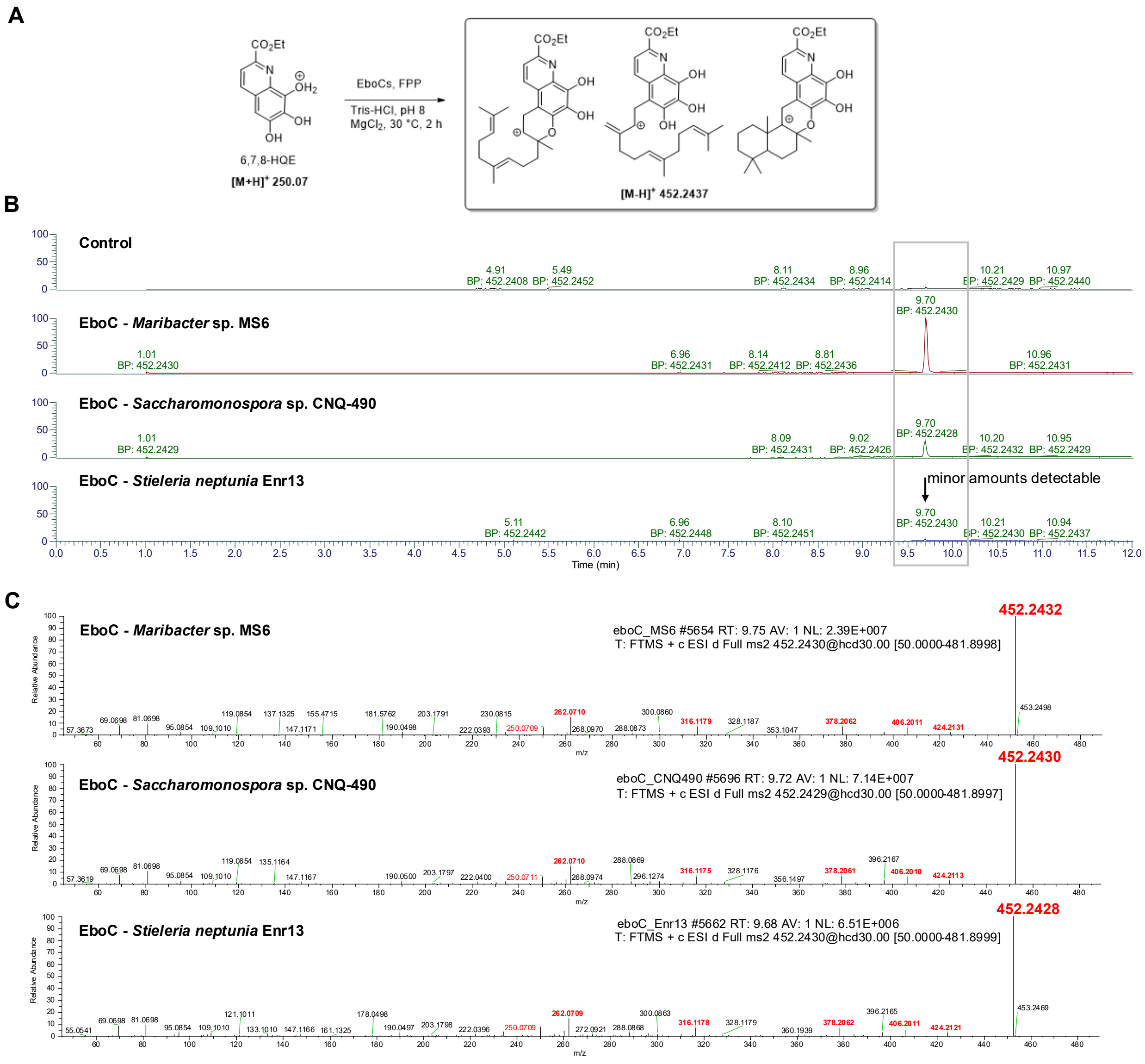


**Fig. S8. Evaluation of EboC homologs.** (**A**) Reaction scheme of enzyme assay using synthetic 6,7,8-trihydroxyquinaldic acid ethyl ester (6,7,8-HQE) as precursor substrate and structures of putative products detectable by LS-MS/MS. (**B**) EIC (*m/z* 452.2437 $[M\text{-}H]^+$) of the farnesylated product in the control sample (empty vector), and heterologously expressed *eboC* homologs of *Maribacter* sp. MS6 (*Bacteroidota*), *Saccharomonospora* sp. CNQ-490 (*Actinomycetota*), and *Stieleria neptunia* Enr13 (*Planctomycetota*). (**C**) $MS^2$ spectrum with detectable fragmentation pattern of *m/z* 452.2437 $[M\text{-}H]^+$ observed from heterologously expressed *eboC* genes from *Maribacter* sp. MS6 (*Bacteroidota*), *Saccharomonospora* sp. CNQ-490 (*Actinomycetota*), and *Stieleria neptunia* Enr13 (*Planctomycetota*).

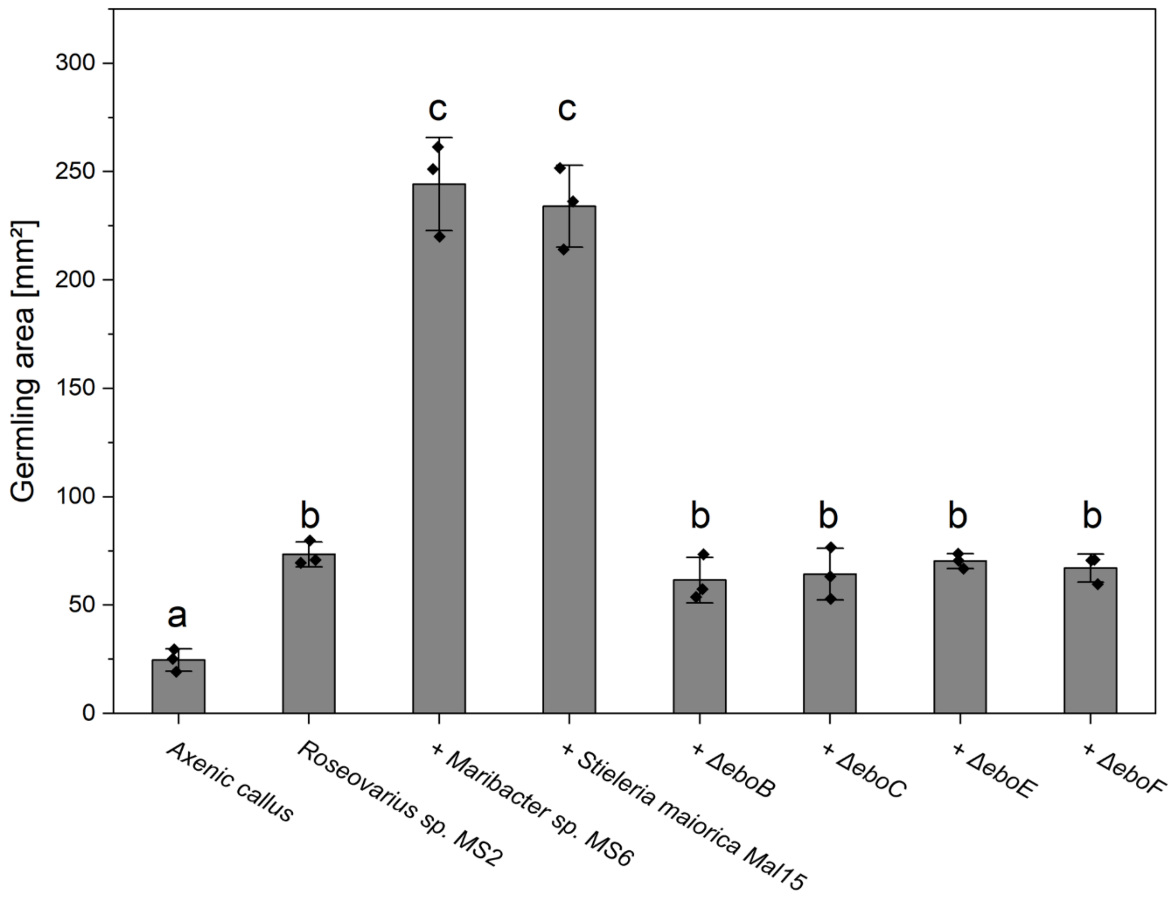


**Fig. S9. Quantification of *Ulva* growth (area) under different bacterial conditions.** Germling area (mm², measured with ImageJ/Fiji software), of *Ulva* after 14 days of cultivation under axenic conditions and upon inoculation with different bacterial strains. Axenic controls show minimal growth and callus-like morphology. Inoculation with *Roseovarius* sp. MS2 results in moderate growth, whereas co-cultivation with *Maribacter* sp. MS6 or *Stieleria maiorica* Mal15 (wildtype and thallusin producer) significantly enhances germling expansion, reflecting complete morphogenesis. In contrast, Δ*ebo* mutants (Δ*eboB*, Δ*eboC*, Δ*eboE*, Δ*eboF*) fail to restore full development and exhibit reduced growth, comparable to that of incomplete morphotypes. Bars represent mean values; individual biological replicates are shown as points, and error bars indicate standard deviation. Different letters denote statistically significant differences between treatments (one-way ANOVA with Tukey's post hoc test, $p < 0.05$).

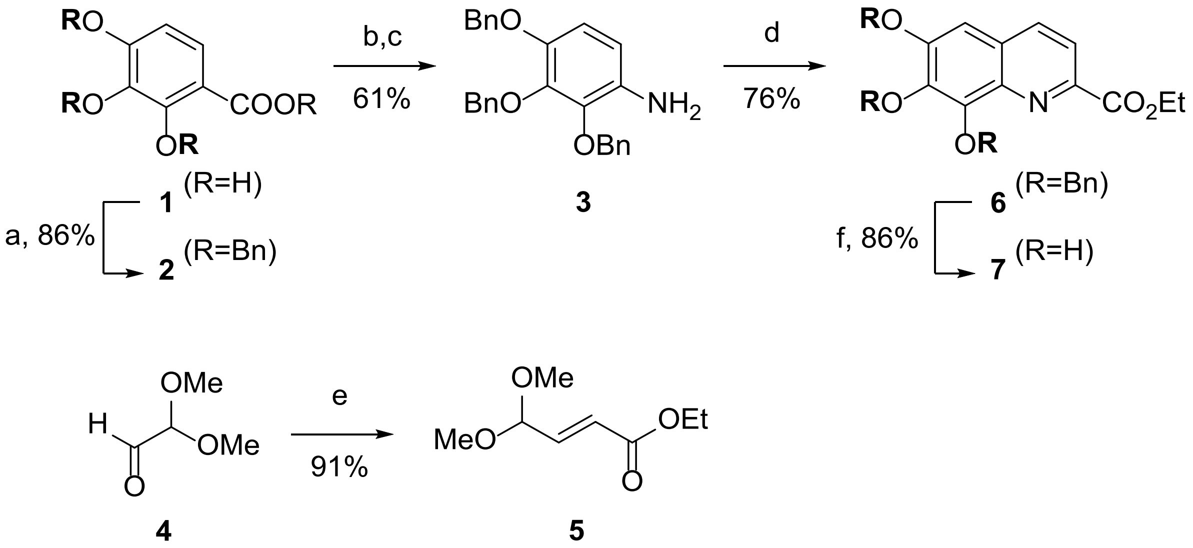


**Fig. S10. Chemical synthesis of the farnesylation precursor 7**. Reagents and conditions: a) BnBr (10.0 eqiv.), $K_2CO_3$, DMF, 80 °C, 16 h; b) LiOH, THF/$H_2O$ 1:1, 70 °C, 12 h; c) diphenylphosphoryl azide (DPPA), triethylamine, 1,4-dioxane, 30 min; then EtOH, KOH, rfx. 2h; d) **5** (1.0 equiv.), iodine (1.0 mol%), trifluoroacetic acid (10 mol%), DMSO, 80 °C, air, 6h; e) triethylphosphonoacetate (1.0 equiv.), $K_2CO_3$ (1.2 equiv.), THF/$H_2O$; f) $H_2$, Pd/C in EtOH/EtOAc 1:1, rt, 1 h.

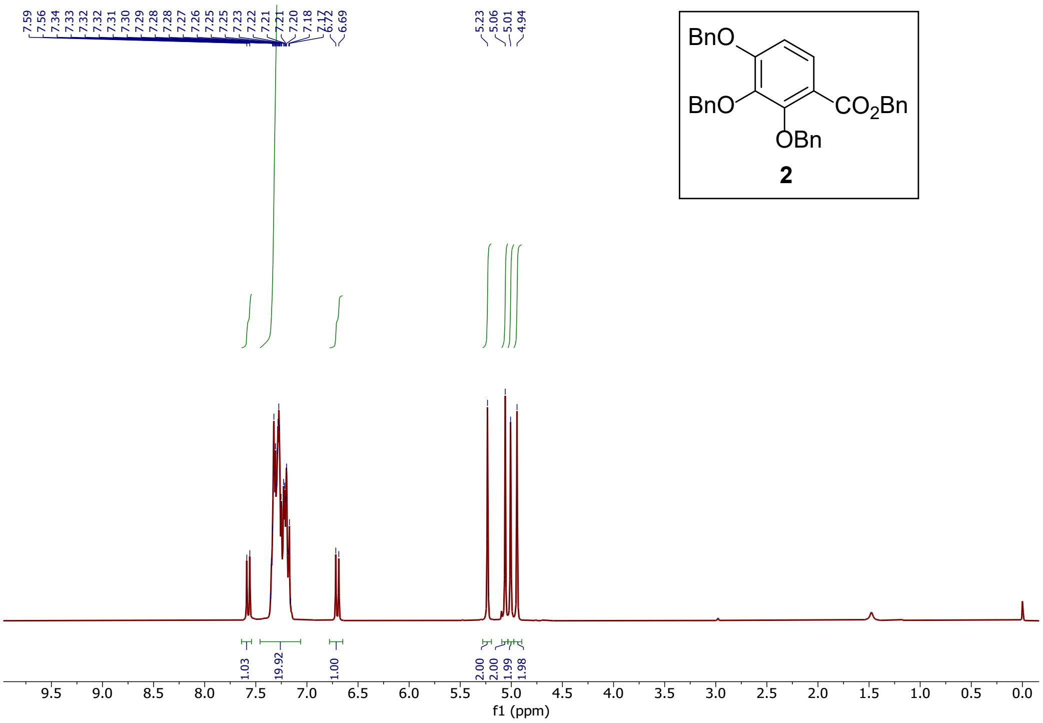


**Fig. S11.** **$^{1}$H NMR (300 MHz, $CDCl_3$) of compound 2**

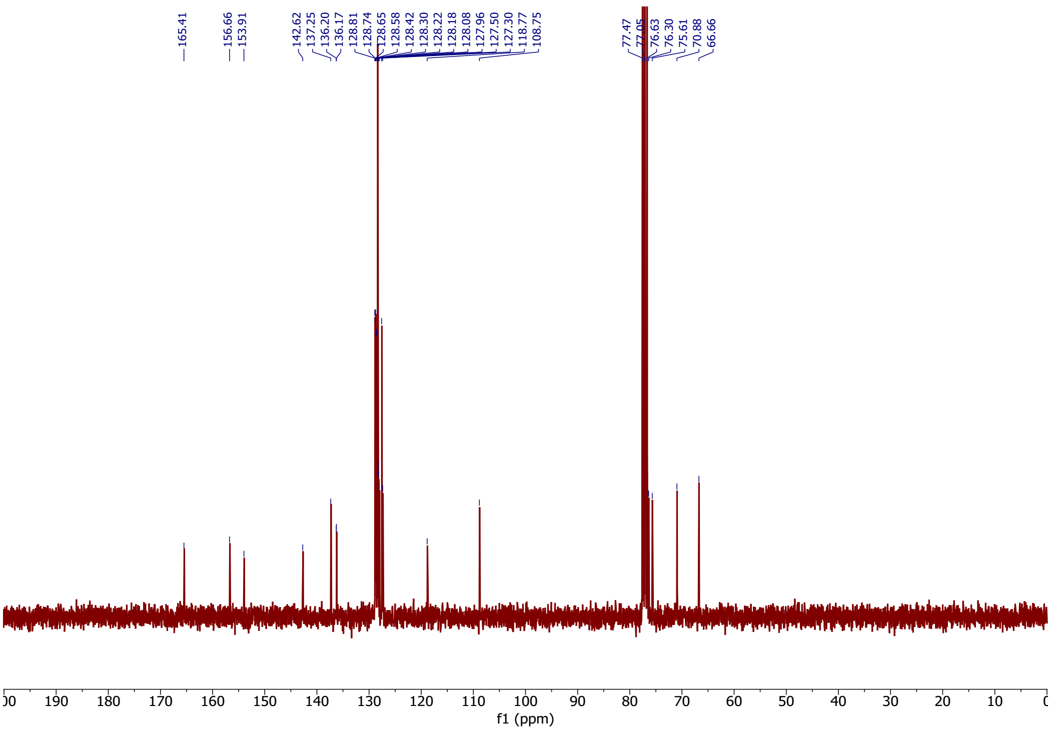


**Fig. S12.** **$^{13}$C NMR (75 MHz, $CDCl_3$) of compound 2**

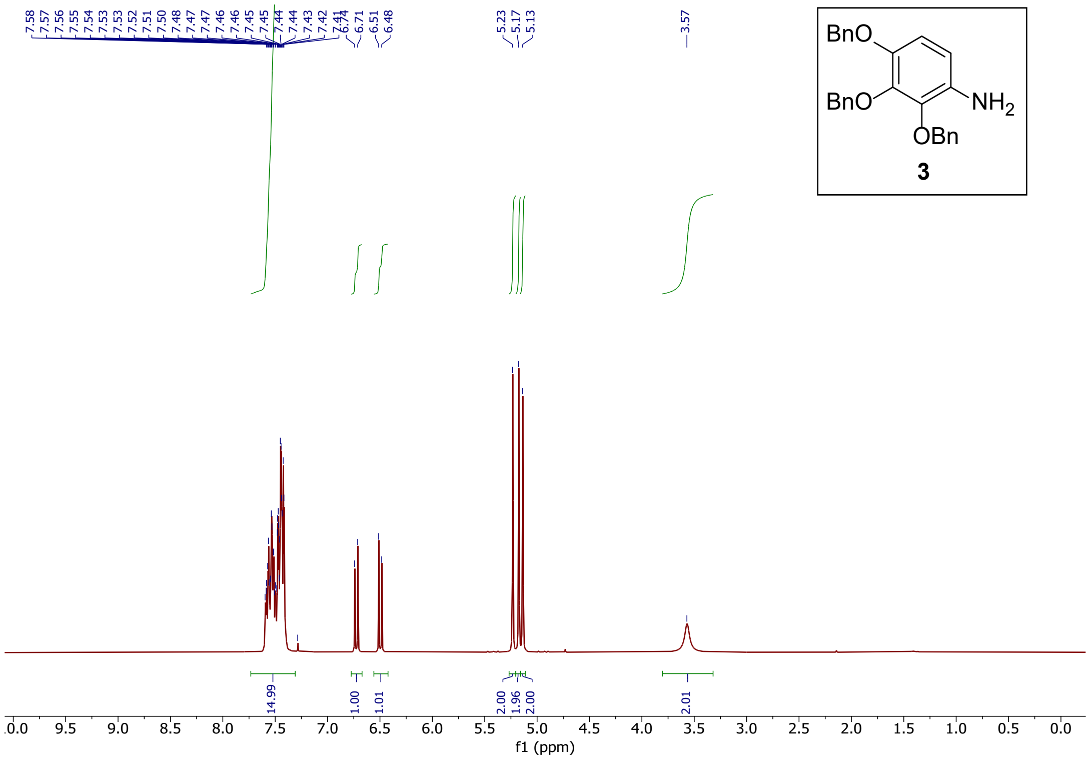


**Fig. S13.** $^{1}$**H NMR (300 MHz,** $CDCl_3$**) of compound 3**

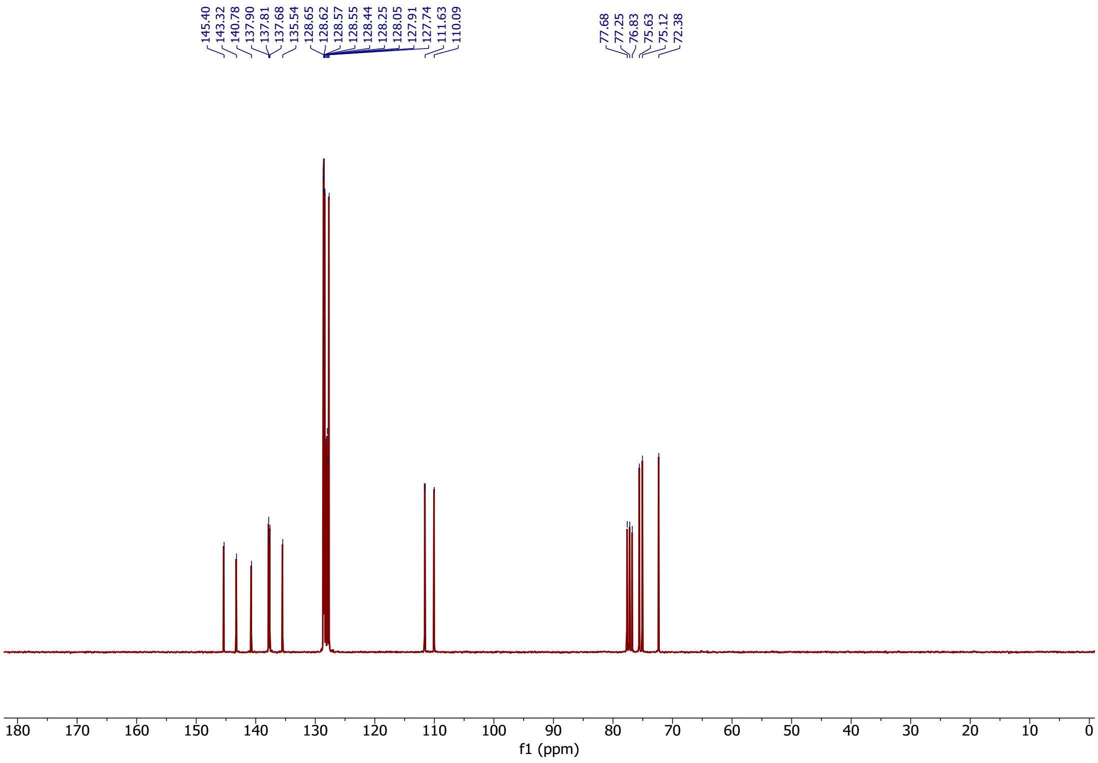


**Fig. S14.** $^{13}$**C NMR (75 MHz,** $CDCl_3$**) of compound 3**

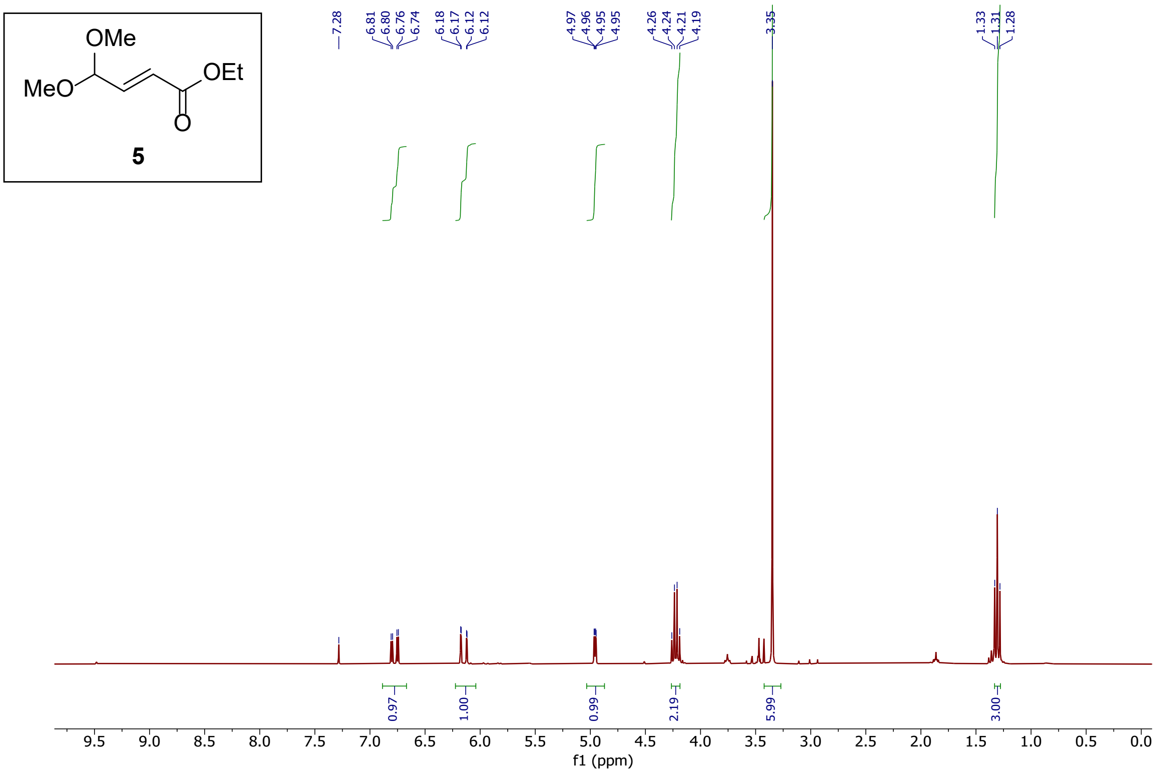


**Fig. S15.** [1]**H NMR (300 MHz, $CDCl_3$) of compound 5**

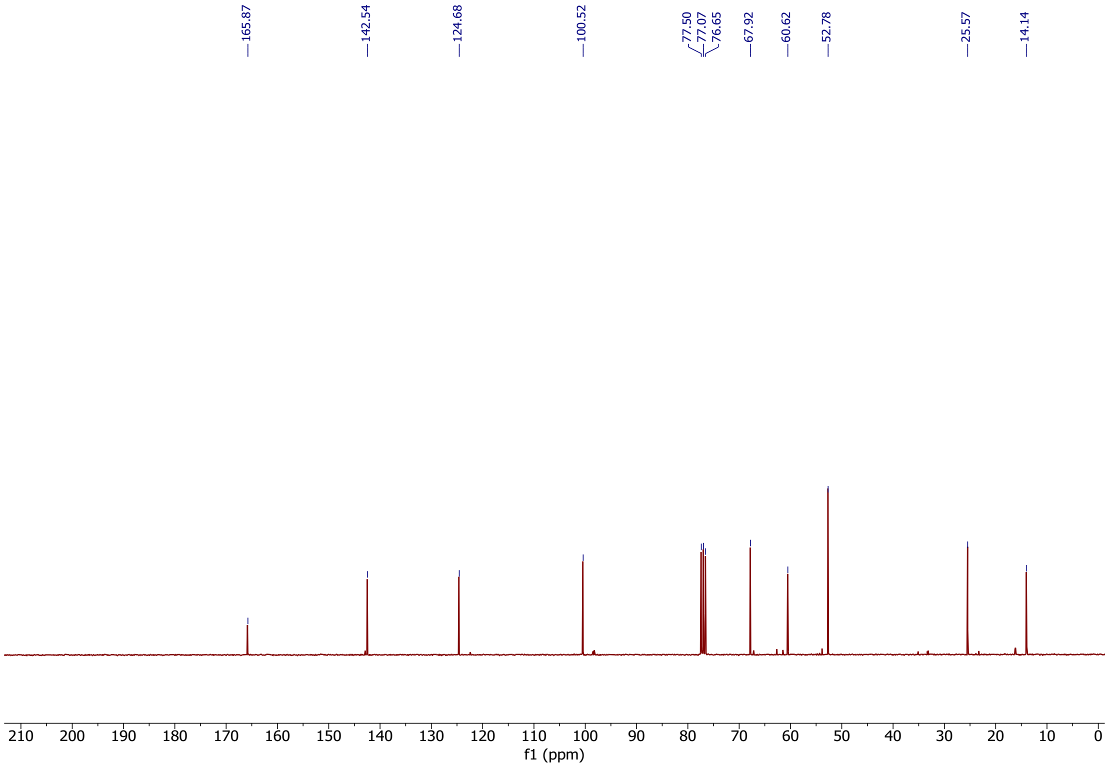


**Fig. S16.** [13]**C NMR (75 MHz, $CDCl_3$) of compound 5**

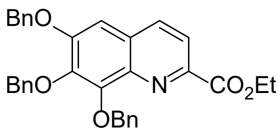


**6**

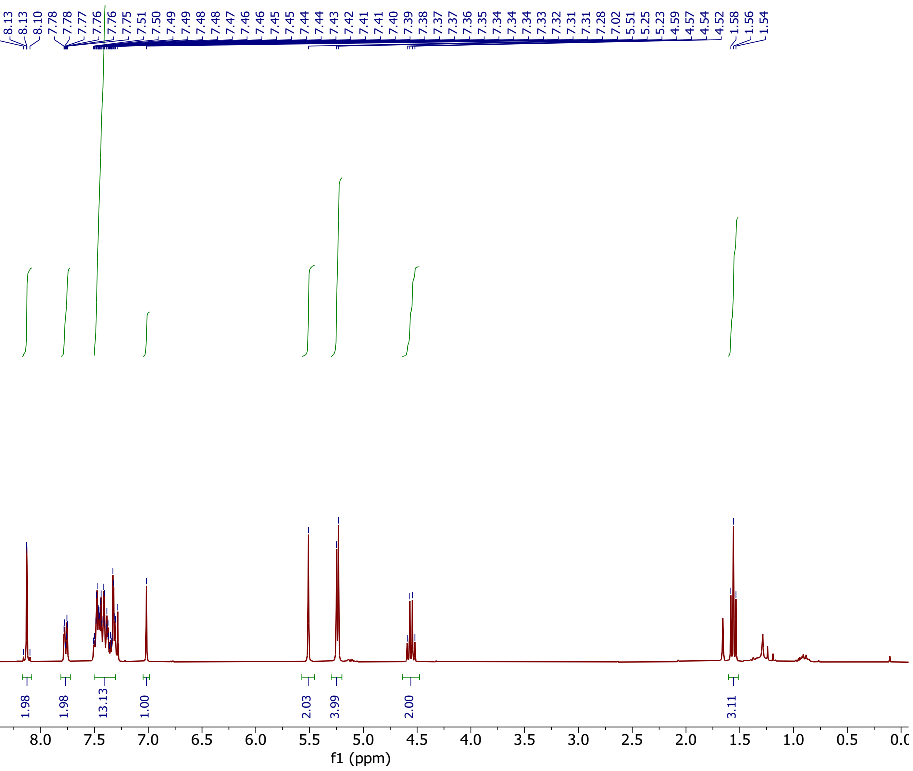


**Fig. S17.** **$^{1}$H NMR (300 MHz, $CDCl_3$) of compound 6**

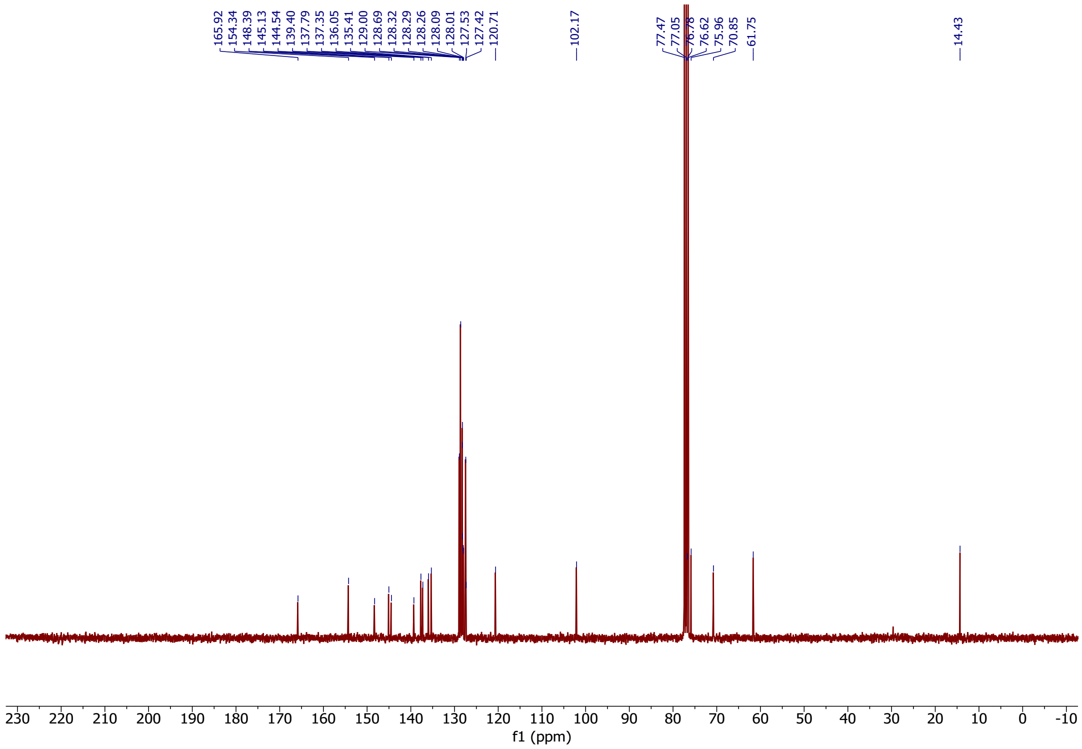


**Fig. S18.** **$^{13}$C NMR (75 MHz, $CDCl_3$) of compound 6**

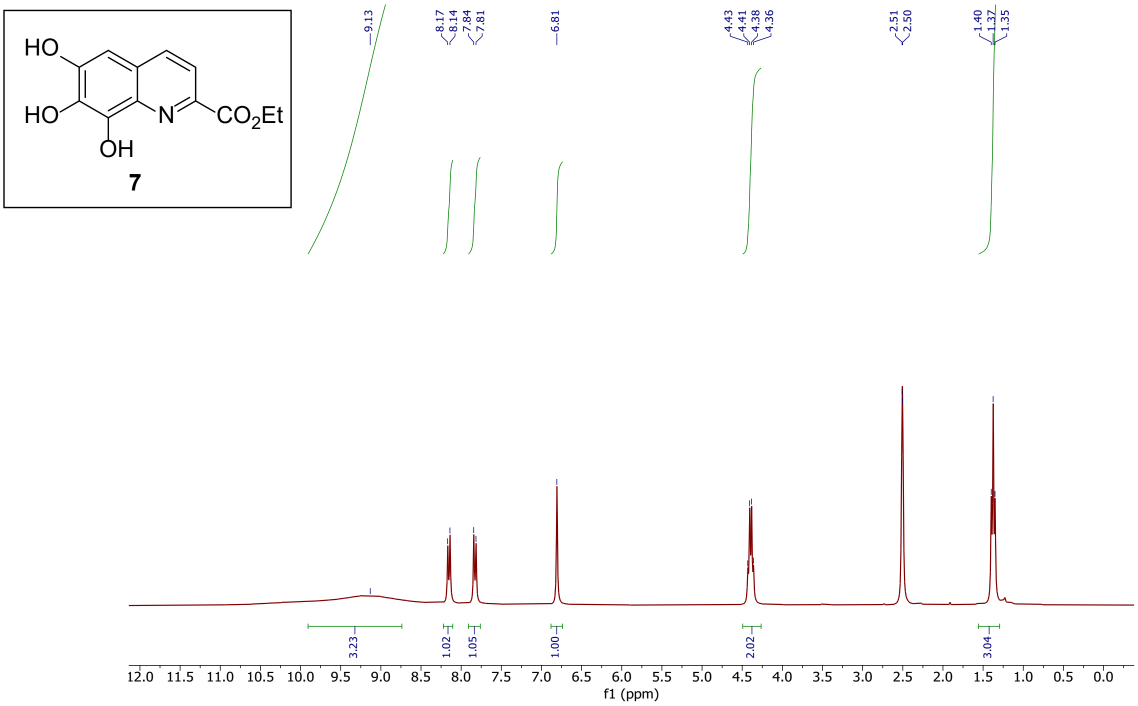


**Fig. S19.** $^{1}$**H NMR (300 MHz, DMSO-D$_6$) of compound 7**

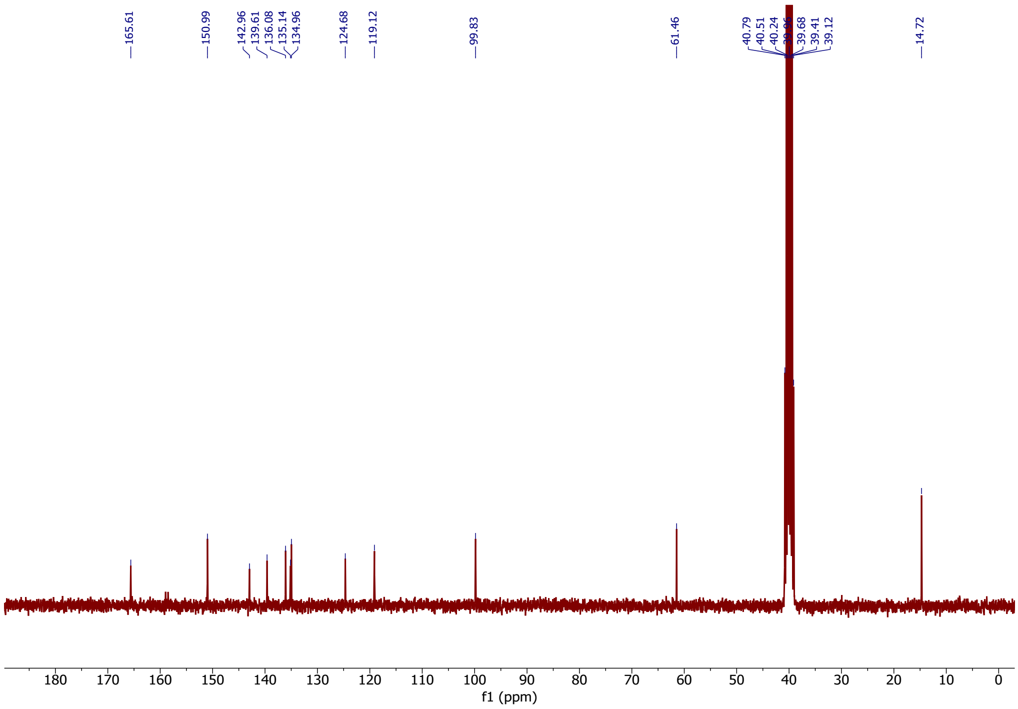


**Fig. S20.** $^{13}$**C NMR (75 MHz, DMSO-D$_6$) of compound 7**

### Supplementary Tables

**Table S1. Morphotypes of *Ulva compressa* in the bacterial morphogenesis bioassay**.

| Morphotype | Description | Inducing Condition | Morphogen Type |
|---|---|---|---|
| **Type I (negative control)** | Callus-like aggregates lacking rhizoids and proper cell walls | Axenic control (no bacteria) | None |
| **Type II** | Enhanced cell division without rhizoid formation | *Roseovarius* sp. MS2 or cytokinin-type activity | Cell-division factor |
| **Type III** | Distinct rhizoid development and cell-wall differentiation | *Maribacter* sp. MS6 or thallusin | Rhizoid-inducing, cell differentiation factor |
| **Type IV (positive control)** | Fully differentiated, morphologically normal thallus | Combination of strains MS2 + MS6 or equivalent isolates | Complete morphogenetic activity |

**Table S2.** RefSeq genome accession numbers of strains tested for thallusin production

| Species name | Strain | Phylum | RefSeq acc. number |
|---|---|---|---|
| *Saccharomonospora* sp. | CNQ-490 | *Actinomycetota* | GCF_000527075.1 |
| *Alteromonas macleodii* | ATCC 27126 | *Pseudomonadota* | GCF_050500325.1 |
| *Roseovarius* sp. | MS2 | *Pseudomonadota* | GCF_051447875.1 |
| *Tranquillimonas alkanivorans* | DSM 19547 | *Pseudomonadota* | GCF_900115595.1 |
| *Saltatorellus ferox* | Poly30 | *Planctomycetota* | GCF_007751475.1 |
| *Rhodopirellula* sp. | UH5 | *Planctomycetota* | GCF_056839225.1 |
| *Rhodopirellula baltica* | SH1 | *Planctomycetota* | GCF_000196115.1 |
| *Roseimaritima multifibrata* | FF011L | *Planctomycetota* | GCF_007741495.1 |
| *Rosistilla ulvae* | EC9 | *Planctomycetota* | GCF_007741475.1 |
| *Rubripirellula lacrimiformis* | K22.7 | *Planctomycetota* | GCF_007741535.1 |
| *Stieleria maiorica* | Mal15 | *Planctomycetota* | GCF_008035925.1 |
| *Stieleria neptunia* | Enr13 | *Planctomycetota* | GCF_007754155.1 |
| *Algoriphagus machipongonensis* | PR1 | *Bacteroidota* | GCF_000166275.1 |
| *Polaribacter dokdonensis* | DSW-5 | *Bacteroidota* | GCF_001280865.1 |
| *Ulvibacter litoralis* | KMM 3912 | *Bacteroidota* | GCF_900102055.1 |
| *Algibacter lectus* | DSM 15365 | *Bacteroidota* | GCF_004368925.1 |
| *Aurantibacter* sp. | UH7 | *Bacteroidota* | GCF_056839165.1 |
| *Flagellimonas zhangzhouensis* | DSM 25030 | *Bacteroidota* | GCF_900106825.1 |
| *Arenibacter palladensis* | DSM 17539 | *Bacteroidota* | GCF_900129275.1 |
| *Pseudozobellia thermophila* | DSM 19858 | *Bacteroidota* | GCF_900141855.1 |
| *Zobellia galactanivorans* | DsiJ | *Bacteroidota* | GCF_000973105.1 |
| *Zobellia uliginosa* | ATCC 14397 | *Bacteroidota* | GCF_900156625.1 |
| *Zobellia russellii* | KMM 3677 | *Bacteroidota* | GCF_018728565.1 |
| *Zobellia amurskyensis* | KMM 3526 | *Bacteroidota* | GCF_009725985.1 |
| *Zobellia laminariae* | KMM 3676 | *Bacteroidota* | GCF_009725995.2 |
| *Maribacter polysiphoniae* | DSM 23514 | *Bacteroidota* | GCF_003148665.1 |
| *Maribacter aestuarii* | JCM 18631 | *Bacteroidota* | GCF_027474845.2 |
| *Maribacter chungangensis* | CAU 1044 | *Bacteroidota* | GCF_042676845.1 |
| *Maribacter* sp. | MS6 | *Bacteroidota* | GCF_056839185.1 |
| *Maribacter sedimenticola* | DSM 19840 | *Bacteroidota* | GCF_900188415.1 |
| *Maribacter arcticus* | DSM 23546 | *Bacteroidota* | GCF_900167935.1 |
| *Maribacter stanieri* | DSM 19891 | *Bacteroidota* | GCF_900112245.1 |
| *Maribacter ulvicola* | DSM 15366 | *Bacteroidota* | GCF_900155985.1 |
| *Maribacter* sp. | BPC-D8 | *Bacteroidota* | GCF_035207705.1 |
| *Maribacter* sp. | UH1 | *Bacteroidota* | GCF_056839145.1 |
| *Nostoc punctiforme* | PCC 73102 | *Cyanobacteriota* | GCF_000020025.1 |
| *Nostoc* sp. | PCC 7120 | *Cyanobacteriota* | GCF_000009705.1 |
| *Synechocystis* sp. | PCC 6803 | *Cyanobacteriota* | GCF_000009725.1 |

**Table S3. Pangenome hits.** COG24 annotation of proteins conserved in thallusin-producing Maribacter spp. but absent in the non-producer *Maribacter aestuarii* JCM 18631. To avoid redundancies only the hits of *Maribacter* sp. UH1 are shown. Blank cells correspond to gene locus IDs to which no COG number could be assigned.

| Gene locus ID | COG24 acc. no. | COG24 function |
|---|---|---|
| 122 | COG0514 | Superfamily II DNA helicase RecQ (RecQ) (PDB:1OYW) |
| 125 | COG2183 | Transcriptional accessory protein Tex/SPT6 (Tex) (PDB:2OCE) |
| 137 | | |
| 303 | COG1988 | Membrane-bound metal-dependent hydrolase YbcI, DUF457 family (YbcI) |
| 327 | COG1228 | Imidazolonepropionase or related amidohydrolase (HutI) (PDB:3OOQ) |
| 460 | COG5279 | Cytokinesis protein 3, contains TGc (transglutaminase/protease-like) domain (CYK3) |
| 461 | | |
| 518 | | |
| 520 | | |
| 589 | COG0624 | Acetylornithine deacetylase/Succinyl-diaminopimelate desuccinylase or related deacylase (ArgE) (PDB:1CG2) |
| 739 | | |
| 745 | COG3564 | Uncharacterized conserved protein, DUF779 family |
| 746 | COG1012 | Acyl-CoA reductase or other NAD-dependent aldehyde dehydrogenase (AdhE) (PDB:1A4S) |
| 747 | COG2207 | AraC-type DNA-binding domain and AraC-containing proteins (AraC) (PDB:1BL0) |
| 782 | COG1028 | NAD(P)-dependent dehydrogenase, short-chain alcohol dehydrogenase family (FabG) (PDB:6L1H) |
| 788 | COG2010\|COG2133 | Cytochrome c, mono- and diheme variants (CccA) (PDB:1A2S)\|Glucose/arabinose dehydrogenase, beta-propeller fold (YliI) (PDB:1C9U) |
| 811 | | |
| 922 | | |
| 1104 | COG0042 | tRNA-dihydrouridine synthase (DusA) (PDB:1VHN) |
| 1135 | COG2197 | DNA-binding response regulator, NarL/FixJ family, contains REC and HTH domains (CitB) (PDB:1A04) |
| 1136 | | |
| 1137 | COG5979 | KinE-related sensor histidine kinase (KinElike) |
| 1140 | | |
| 1141 | COG3279 | DNA-binding response regulator, LytR/AlgR family (LytT) (PDB:3BS1) |

| | | |
|---|---|---|
| 1147 | COG0463\|COG0745 | Glycosyltransferase involved in cell wall bisynthesis (WcaA) (PDB:5MLZ)\|DNA-binding response regulator, OmpR family, contains REC and winged-helix (wHTH) domain (OmpR) (PDB:1XHF) |
| 1153 | COG0463\|COG0745 | Glycosyltransferase involved in cell wall bisynthesis (WcaA) (PDB:5MLZ)\|DNA-binding response regulator, OmpR family, contains REC and winged-helix (wHTH) domain (OmpR) (PDB:1XHF) |
| 1207 | | |
| 1292 | | |
| 1375 | COG2905 | Signal-transduction protein containing cAMP-binding, CBS, and nucleotidyltransferase domains |
| 1382 | COG1169 | Isochorismate synthase EntC (MenF) (PDB:5JXZ) |
| 1400 | | |
| 1418 | COG2143 | Thioredoxin-related protein SoxW (SoxW) (PDB:4FYB) |
| 1527 | COG1171 | Threonine deaminase (IlvA) (PDB:3IAU) |
| 1685 | COG3204 | Uncharacterized Ca-binding beta-propeller protein YjiK (YjiK) (PDB:3QQZ) |
| 1704 | | |
| 1719 | | |
| 1868 | | |
| 1889 | COG1076 | DnaJ domain-containing protein (DjlA) (PUBMED:15489435;11106641) |
| 1890 | COG0513 | Superfamily II DNA and RNA helicase (SrmB) (PDB:3RRM) |
| 1891 | COG1028 | NAD(P)-dependent dehydrogenase, short-chain alcohol dehydrogenase family (FabG) (PDB:6L1H) |
| 1914 | | |
| 1955 | COG2510 | Riboflavin transporter RibN, EamA domain (RibN) (PUBMED:23935051) |
| 1960 | COG0702 | Uncharacterized conserved protein YbjT, contains NAD(P)-binding and DUF2867 domains (YbjT) (PDB:5L3Z) |
| 1988 | COG0352 | Thiamine monophosphate synthase (ThiE) (PDB:1G4T) (PUBMED:19060138) |
| 1993 | COG0352 | Thiamine monophosphate synthase (ThiE) (PDB:1G4T) (PUBMED:19060138) |
| 2018 | COG0545 | FKBP-type peptidyl-prolyl cis-trans isomerase (FkpA) (PDB:1A7X) |
| 2019 | | |
| 2020 | COG0526 | Thiol-disulfide isomerase or thioredoxin (TrxA) (PDB:1XFL) |
| 2021 | COG1047 | Peptidyl-prolyl cis-trans isomerase, FKBP type (SlpA) (PDB:2M2A) |
| 2078 | COG0697 | Amino acid export permease, drug/metabolite transporter (DMT) superfamily (EamA) (PDB:5I20) (PUBMED:11432728;17784858;35495724;35598887) |
| 2185 | | |
| 2200 | | |

| | | |
|---|---|---|
| 2239 | COG1733 | DNA-binding transcriptional regulator, HxlR family (HxlR) (PDB:1YYV) |
| 2256 | COG3605 | Signal transduction protein containing GAF and PtsI domains (PtsP) |
| 2403 | | |
| 2421 | COG2608 | Copper chaperone CopZ (CopZ) (PDB:1AFI) |
| 2451 | COG1721 | Uncharacterized membrane-anchored protein with extracellular vWFA and Ig-like domains, component of a predicted archaeal secretion system (MMP0362) (PUBMED:25583072) |
| 2453 | | |
| 2454 | COG0312 | Zn-dependent protease or N-deacetylase, PmbA/TldD/TldE family (TldD) (PDB:3TV9) (PUBMED:12029038;28943336;36155963) |
| 2455 | COG0312 | Zn-dependent protease or N-deacetylase, PmbA/TldD/TldE family (TldD) (PDB:3TV9) (PUBMED:12029038;28943336;36155963) |
| 2516 | COG4232 | Thiol:disulfide interchange protein DsbD (DsbD) (PDB:1JPE) |
| 2608(*eboF*) | COG1524 | c-di-AMP phosphodiesterase AtaC or nucleotide pyrophosphatase, AlkP superfamily (AtaC) (PDB:1EI6) (PUBMED:32188788) |
| 2609(*eboE*) | | |
| 2610(*eboD*) | COG0337 | 3-dehydroquinate synthetase (AroB) (PDB:3ZOK) |
| 2611(*eboC*) | COG0382 | 4-hydroxybenzoate polyprenyltransferase (UbiA) (PDB:4OD4) (PUBMED:28830929) |
| 2612(*eboB*) | COG1099 | Predicted metal-dependent hydrolase, TIM-barrel fold (PDB:3GUW) |
| 2613(*eboA*) | | |
| 2784 | COG2197 | DNA-binding response regulator, NarL/FixJ family, contains REC and HTH domains (CitB) (PDB:1A04) |
| 2828 | COG3264 | Small-conductance mechanosensitive channel MscK (MscK) (PDB:7UW5) (PUBMED:17493135;36371466) |
| 2829 | | |
| 2830 | COG1595 | DNA-directed RNA polymerase specialized sigma subunit, sigma24 family (RpoE) (PDB:2Q1Z) |
| 2944 | | |
| 3062 | | |
| 3153 | COG0784 | CheY-like REC (receiver) domain, includes chemotaxis protein CheY and sporulation regulator Spo0F (CheY) (PDB:6QRJ) |
| 3193 | COG4886 | Type III secretion system effector YopM, contains leucine-rich repeats (YopM) (PDB:4OW2) |
| 3196 | | |
| 3220 | | |
| 3228 | COG0772 | Peptodoglycan polymerase FtsW/RodA/SpoVE (FtsW) (PDB:6BAR) (PUBMED:30692671) |
| 3278 | COG0500 | SAM-dependent methyltransferase SmtA (CmoB moved to COG2228) (SmtA) (PDB:5DNK) (PUBMED:8566713) |
| 3379 | COG1929 | Glycerate kinase (GlxK) (PDB:1TO6) |
| 3437 | COG0697 | Amino acid export permease, drug/metabolite transporter (DMT) superfamily (EamA) (PDB:5I20) (PUBMED:11432728;17784858;35495724;35598887) |

| | | |
|---|---|---|
| 3458 | COG0784 | CheY-like REC (receiver) domain, includes chemotaxis protein CheY and sporulation regulator Spo0F (CheY) (PDB:6QRJ) |
| 3465 | | |
| 3516 | COG0824 | Acyl-CoA thioesterase FadM (FadM) (PDB:2FUJ) |
| 3520 | COG2761 | Predicted dithiol-disulfide isomerase, DsbA/YjbH family (virulence, stress resistance) (FrnE) (PDB:6GHB) |
| 3754 | COG3129 | 23S rRNA A1618 N6-methylase RlmF (RlmF) (PDB:2H00) (PUBMED:18021804) |
| 3755 | | |
| 3804 | | |
| 3827 | COG0356 | FoF1-type ATP synthase, membrane subunit a (AtpB) (PDB:1C17_M) (PUBMED:22931285) |
| 3882 | COG0673 | Predicted dehydrogenase (MviM) (PDB:3UUW) |
| 3945 | | |

### Table S4. Details on materials and tools used for genome sequencing

| Strain name | UH1 | UH5 | UH7 |
|---|---|---|---|
| **DNA Extraction** | Wizard® HMW DNA Extraction Kit (Promega) | Wizard® HMW DNA Extraction Kit (Promega) | Whole genome amplication from early-exponential cultures with REPLI-g Single Cell Kit (Qiagen) |
| **ONT Sequencing Kit** | SQK-NBD114.24 | SQK-NBD114.24 | SQK-NBD114.24 |
| **ONT Flow cell** | FLO-MIN114 | FLO-MIN114 | FLO-MIN114 |
| **Bascaller** | Dorado 1.3.1 | Dorado 1.1.1 | Dorado 1.0.0 |
| **Basecalling_model** | dna_r10.4.1_e8.2_400bps_sup@v5.2.0 | dna_r10.4.1_e8.2_400bps_sup@v5.2.0 | dna_r10.4.1_e8.2_400bps_sup@v5.2.0 |
| **Demultiplexing** | Dorado 1.3.1 | Dorado 1.1.1 | Dorado 1.0.0 |
| **Q10 filtering** | Dorado 1.3.1 | Dorado 1.1.1 | Dorado 1.0.0 |
| **Read QC long reads** | NanoPlot 1.46.1 | NanoPlot 1.46.1 | NanoPlot 1.44.1 |
| **Assembly** | Flye 2.9.6 (--nano-hq -m 2000) | Flye 2.9.6 (--nano-hq) | Flye 2.9.6 (--nano-hq) |
| **Long-read Polishing** | Medaka 1.7.2 | Medaka 1.7.2 | Medaka 1.7.2 |
| **Medaka Model** | r1041_e82_400bps_sup_g615 | r1041_e82_400bps_sup_g615 | r1041_e82_400bps_sup_g615 |
| **Read QC short reads** | FastQC 0.74 | FastQC 0.74 | FastQC 0.74 |
| **Mappig of short reads** | BWA-MEM2 2.3 | BWA-MEM2 2.2.1 | BWA-MEM2 2.2.1 |
| **Short-read polishing** | Pilon 1.20.1 | Pilon 1.20.1 | Pilon 1.20.1 |
| **Evaluation of completeness** | BUSCO 5.8.0 (--lineage_dataset 'flavobacteriia' --metaeuk) | BUSCO 5.8.0 (--lineage_dataset 'planctomycetes_odb10' --metaeuk) | BUSCO 5.8.0 (--lineage_dataset 'flavobacteriales_odb10' --metaeuk) |
| **Annotation for rotation** | Prokka 1.14.6 | Prokka 1.14.6 | Prokka 1.14.6 |

**Table S5. Strains and plasmids are used in this study.**

| Strain or plasmid | Purpose or relevant characteristics | Source or reference |
|---|---|---|
| **Strains** | | |
| ***Escherichia coli* DH5α** | general purpose cloning strain | Thermo Fisher |
| ***E. coli* TOP10** | general purpose cloning strain | Thermo Fisher |
| ***E. coli* BL21(DE3)** | strain for heterologous gene expression | Thermo Fisher |
| ***Stieleria maiorica* Mal15** | type strain DSM 100215 | (*1*) |
| ***S. maiorica* Mal15 Δ*eboB*** | derivative in which *eboB* (Mal15_37820) is replaced by the *cat* gene | This study |
| ***S. maiorica* Mal15 Δ*eboC*** | derivative in which *eboC* (Mal15_01440) is replaced by the *cat* gene | This study |
| ***S. maiorica* Mal15 Δ*eboE*** | derivative in which *eboE* (Mal15_49730) is replaced by the *cat* gene | This study |
| ***S. maiorica* Mal15 Δ*eboF*** | derivative in which *eboF* (Mal15_03990) is replaced by the *cat* gene | This study |
| **Plasmids** | | |
| **pDEL(cat)** | *E. coli* vector (ColE1 ori$_{Ec}$), *cm*$^R$ | (*2*) |
| **pDEL(cat)_Mal15_37820-updown** | pDEL(cat) derivative harbouring the homology arms flanking the gene Mal15_37820 | This study |
| **pDEL(cat)_Mal15_01440-updown** | pDEL(cat)derivative harbouring the homology arms flanking the gene Mal15_01440 | This study |
| **pDEL(cat)_Mal15_49730-updown** | pDEL(cat) derivative harbouring the homology arms flanking the gene Mal15_49730 | This study |
| **pDEL(cat)_Mal15_03990-updown** | pDEL(cat) derivative harbouring the homology arms flanking the gene Mal15_03990 | This study |
| **pET28a** | *E. coli* vector (ColE1 ori$_{Ec}$), f1-ori, T7 promoter and terminator, *lacI*, *kan*$^R$ | Novagen (Merck Millipore) |
| **pAGB026** | pET28a derivative harboring *eboC* from *Maribacter* sp. MS6 (codon-optimized for *E. coli*) | This study |
| **pAGB031** | pET28a derivative harboring *eboC* from *Stieleria neptunia* Enr13 (codon-optimized for *E. coli*) | This study |
| **pAGB032** | pET28a derivative harboring *eboC* from *Saccharomonospora sp.* CNQ490 (codon-optimized for *E. coli*) | This study |
| ***cm*$^r$*:* chloramphenicol resistance*,* *kan*$^r$: kanamycin resistance** | | |

**Table S6. Oligonucleotides are used in this study. Restriction sites used for cloning are underlined.**

| Primer name | Sequence (5'-3') | Site |
|---|---|---|
| **XhoI-eboB_Sm-up-s** | TCT**CTCGAG**GTCCGGTATTGCAACCACCAGATTCGGACTC | XhoI |
| **KpnI-eboB_Sm-up-as** | TCT**GGTACC**GATGTGCGGGTCGATGTAGTCCATACGTCATCAATCAAG | KpnI |
| **XbaI-eboB_Sm-down-s** | TAA**TCTAGA**TTTAATTTCACCCCACGGGGTTAGGGCTTCCGTCAATTC | XbaI |
| **HindIII-eboB_Sm-down-as** | TCT**AAGCTT**CGCGTCAGGTAGCTATGCAAGGCGGCGAC | HindIII |
| **check-eboB_Sm-s** | GGGTGATCACCTTGTTGATCTACCACGCCTCGATTG | -- |
| **check-eboB_Sm-as** | TAGGGTGATAAAAGCTGTCTTCCAATTTCAACAGATTGCGACGGGTG | -- |
| **KpnI-eboC_Sm-up-s** | CTC**GGTACC**ATGCTCAGTCGGGGACAGACCCCGAG | KpnI |
| **NheI-eboC_Sm-up-as** | CTC**GCTAGC**GTTGGGCAGACGAACCAGTTGCATCCAG | NheI |
| **XbaI-eboC_Sm-down-s** | CTC**TCTAGA**GCGACGCGATTGCGGGTGACGTGAGATG | XbaI |
| **HindIII-eboC_Sm-down-as** | CTC**AAGCTT**CCCTTCGTCGGAACCGTGGGATGCTG | HindIII |
| **check-eboC_Sm-s** | AGACCCCGATCCGTTGGGGACAGAC | -- |
| **check-eboC_Sm-as** | AAGGTTCAATCCGACGAAGATCGTCTCAATCGTTACGAC | -- |
| **XhoI-eboE_Sm-up-s** | TCT**CTCGAG**AAACCACTGCGCGATCTCATAGATGTAGTTCTTGTCGTCTTCAAAGAATTCCTTG | XhoI |
| **NheI-eboE_Sm-up-as** | TCT**GCTAGC**TGGGTCTGAATCGTAGATGTTCAATGTGCTAACATGACGGAG | NheI |
| **XbaI-eboE_Sm-down-s** | TAA**TCTAGA**CTGGCCGATCAACTCGCGGGATAGGCTTG | XbaI |
| **HindIII-eboE_Sm-down-as** | TCT**AAGCTT**AGCCCCCAGATTGTCGTGCCCAAAGCCAC | HindIII |
| **check-eboE_Sm-s** | CCAACGTGAACTCACCGCGTCCCAGGAACGAC | -- |
| **check-eboE_Sm-as** | TCGCAGCGATTTCGACATCCAAAGCCGATCTC | -- |
| **XhoI-eboF_Sm-up-s** | TCT**CTCGAG**GGATCTTTAGCGTCGTCTGCCGAAGTGCCAAG | XhoI |
| **KpnI-eboF_Sm-up-as** | TCT**GGTACC**ATTGATCACACAAACCCGATCCATTCACTCAGGTCCGCTTC | KpnI |
| **XbaI-eboF_Sm-down-s** | TAA**TCTAGA**CGGTCGCTGCTACGCGGCCGTTGAAC | XbaI |
| **HindIII-eboF_Sm-down-as** | TCT**AAGCTT**CGACGGCGACCAACGTTCCATCGGGGGAAAC | HindIII |
| **check-eboF_Sm-s** | GGTGCTGGGGATTCCGTTGTGCTGG | -- |
| **check-eboF_Sm-as** | CGATTCCCGTCACCGGATCGCGGTC | -- |
| **eboB_Sm-in-s** | GGGCTGCGTGGCAATGAGCGAACCTG | -- |
| **eboB_Sm-in-as** | CGAGCACGTCGGTGATCGTGTGCTCTTCGAC | -- |
| **eboC_Sm_in-s** | GATGTTTTCGATTTGGAGAAAGACAAGGCGGAGCGGAAAATG | -- |
| **eboC_Sm_in-as** | ATCAACCCGATCAACATTCCGAACTGACCCGCCGAC | -- |
| **eboE_Sm-in-s** | TCGATCCGCGGCAATCTGCTGGAGTATG | -- |
| **eboE_Sm-in-as** | GGTCGATTCGAACAGCTGATCCAGGAACCCTGACAATTTAC | -- |
| **eboF_Sm-in-s** | AACTTCATGCCGTGGACGAGCCCGTTG | -- |
| **eboF_Sm-in-as** | TGTCATCGGCGGCGGACAAGATGGTTTG | -- |

**Table S7. Solvents, reagents, and isotopically labeled compounds are listed in alphabetical order.**

| Compound | Labeling | Purity (%) | Role | Supplier |
|---|---|---|---|---|
| **Anthranilic acid** | $^{15}N$ | 98 | Precursor | Cambridge Isotope Laboratories, US |
| **L-aspartic acid** | $^{15}N$ | 98 | Key precursor | Cambridge Isotope Laboratories, US |
| **L-aspartic acid** | 4-$^{13}C$ | 99 | Key precursor | Cambridge Isotope Laboratories, US |
| **L-aspartic acid** | $^{13}C_4{}^{15}N$ | 98 | Key precursor | Cambridge Isotope Laboratories, US |
| **Erythrose** | 1-$^{13}C$ | 98 | Carbon precursor | Cambridge Isotope Laboratories, US |
| **L-glutamic acid** | $^{13}C_5{}^{15}N$ | 98 | Indirect precursor | Cambridge Isotope Laboratories, US |
| **L-tryptophan** | $^{13}C_{11}{}^{15}N_2$ | 99 | Indirect precursor | Cambridge Isotope Laboratories, US |
| **Methanol** | — | HPLC grade | Solvent | VWR, Germany |
| **KOH** | — | — | Base | VWR, Germany |
| **DMSO** | — | — | Solvent | Carl Roth, Germany |
| **Marine broth** | — | — | Growth medium | Carl Roth, Germany |
| **Iodomethane** | — | — | Derivatization reagent | Sigma-Aldrich, Germany |
| **Water** | — | — | Purified solvent | Th. Geyer, Germany |
| **Formic acid** | — | ≥ 99 | Mobile phase additive | Scientific Thermo Fisher, Germany |
| **UHPLC solvents** | — | UHPLC grade | Solvent | Th. Geyer, Germany |
